# Measurement of stretch-evoked brainstem function using fMRI

**DOI:** 10.1101/2020.06.19.161315

**Authors:** Andrea Zonnino, Andria J Farrens, David Ress, Fabrizio Sergi

## Abstract

Knowledge on the organization of motor function in the reticulospinal tract (RST) is limited by the lack of methods for measuring RST function in humans. Behavioral studies suggest the involvement of the RST in long latency responses (LLRs). LLRs, elicited by precisely controlled perturbations, can therefore act as a viable paradigm to measure motor-related RST activity using functional Magnetic Resonance Imaging (fMRI).

Here we present StretchfMRI, a novel technique developed to study RST function associated with LLRs. StretchfMRI combines robotic perturbations with electromyography and fMRI to simultaneously quantify muscular and neural activity during stretch-evoked LLRs without loss of reliability. Using StretchfMRI, we established the muscle-specific organization of LLR activity in the brainstem. The observed organization is partially consistent with animal models, with activity primarily in the ipsilateral medulla for flexors and in the contralateral pons for extensors, but also include other areas, such as the midbrain and bilateral pontomedullary contributions.

## Introduction

Multiple secondary pathways are known to participate, together with the corticospinal tract, in controlling voluntary motor actions^1^. Among these secondary pathways, the reticulospinal tract (RST) is especially important for its involvement in locomotion^2^, maintenance of posture^3^, reaching^4^ and grasping^5^. The RST is composed of two different pathways that originate from the Reticular Formation (RF), a region in the brainstem composed of a constellation of nuclei^6, 7^. Evidence from non-human primate studies suggests a possible organization of motor function in the RF, where excitatory signals are sent to ipsilateral flexors and contralateral extensors, and inhibitory signals are sent to contralateral flexors and ipsilateral extensors^8–10^.

A complete understanding of how motor functions are organized in human RF however, is still lacking. Due to the small size of the brainstem and its location deep in the cranium, non-invasive measurements are currently achievable using neuroimaging. Unfortunately, the between-subject and between-task variability associated with experiments involving voluntary responses, combined with the relatively low signal-to-noise ratio of functional neuroimaging of the brainstem^11^, make these investigations very challenging.

Recent studies, however, suggest that the RF might be actively involved in the generation of long latency responses (LLRs), a stereotypical response evoked in muscles after an unexpected perturbation displaces one or more joints, thus stretching a set of muscles^12^. Because, LLRs are “semi-reflexive” responses, they are less affected by confounds such as individual subject skill and task performance, resulting in smaller between-subject variability than is found for voluntary motor tasks. As such, precisely evoked LLRs may be a means to reliably stimulate the RST to enable the direct measurement of RF activity using neuroimaging.

Here, we present StretchfMRI, a novel technique that we have developed to study the brainstem correlates of LLRs *in-vivo* in humans. StretchfMRI combines robotic perturbations with electromyography (EMG) and functional Magnetic Resonance Imaging (fMRI) to provide simultaneous recording of neural and muscular activity associated with LLRs. In this paper, we demonstrate that StretchfMRI enables the reliable quantification of both EMG and fMRI associated with LLRs, and establish muscle-specific representation of LLR activity in the brainstem. Additionally, we present an exploratory analysis to determine cortical areas associated with LLR activity for flexors and extensors.

## Results

Healthy individuals were exposed to two experiments after providing informed consent under approval by the Institutional Review Board of the University of Delaware. One experiment (Exp 1, n_1_ = 12 subjects) aimed to evaluate the reliability of EMG measurements collected during fMRI using a four-session design. Experiments were conducted outside (OUT) and inside (IN) the scanner, with two IN sessions (IN_1_ and IN_2_) temporally interleaved between two OUT sessions (OUT_1_ and OUT_2_).

The second experiment (Exp 2, n_2_ = 18 subjects) included two repeated fMRI sessions, and was conducted to to test the hypothesis that the BOLD signal in any location in the brainstem was significantly associated with LLRs for flexors and/or extensor muscles under stretch, and to quantify the reliability of the two repeated fMRI experiments (see Online Methods).

### StretchfMRI

To enable reliable measurement of stretch-evoked muscle responses during fMRI, we combined a novel MRI-compatible robot with custom EMG acquisition and processing methods, and used a modified fMRI sequence including a 225 ms silent window after every acquisition volume, during which stretch-evoked responses were measured (Fig. 1). The robot was used to apply Ramp-and-Hold perturbations at different velocities to condition LLRs of two muscles (Flexor Carpi Radialis - FCR, and Extensor Carpi Ulnaris - ECU). Muscle response was quantified using a custom electrode set that included co-located measurement and reference electrodes. Use of the custom electrode set allowed to simultaneously record both muscle signal and motion artifact induced by the unavoidable movement of the electrode set in the scanner. The measured signals were filtered using a pipeline that included adaptive noise cancellation to estimate and remove the signal related to motion artifacts (non-linearly related to the measurement from the reference electrode), and obtain clean measurements of EMG.

**Figure 1.**
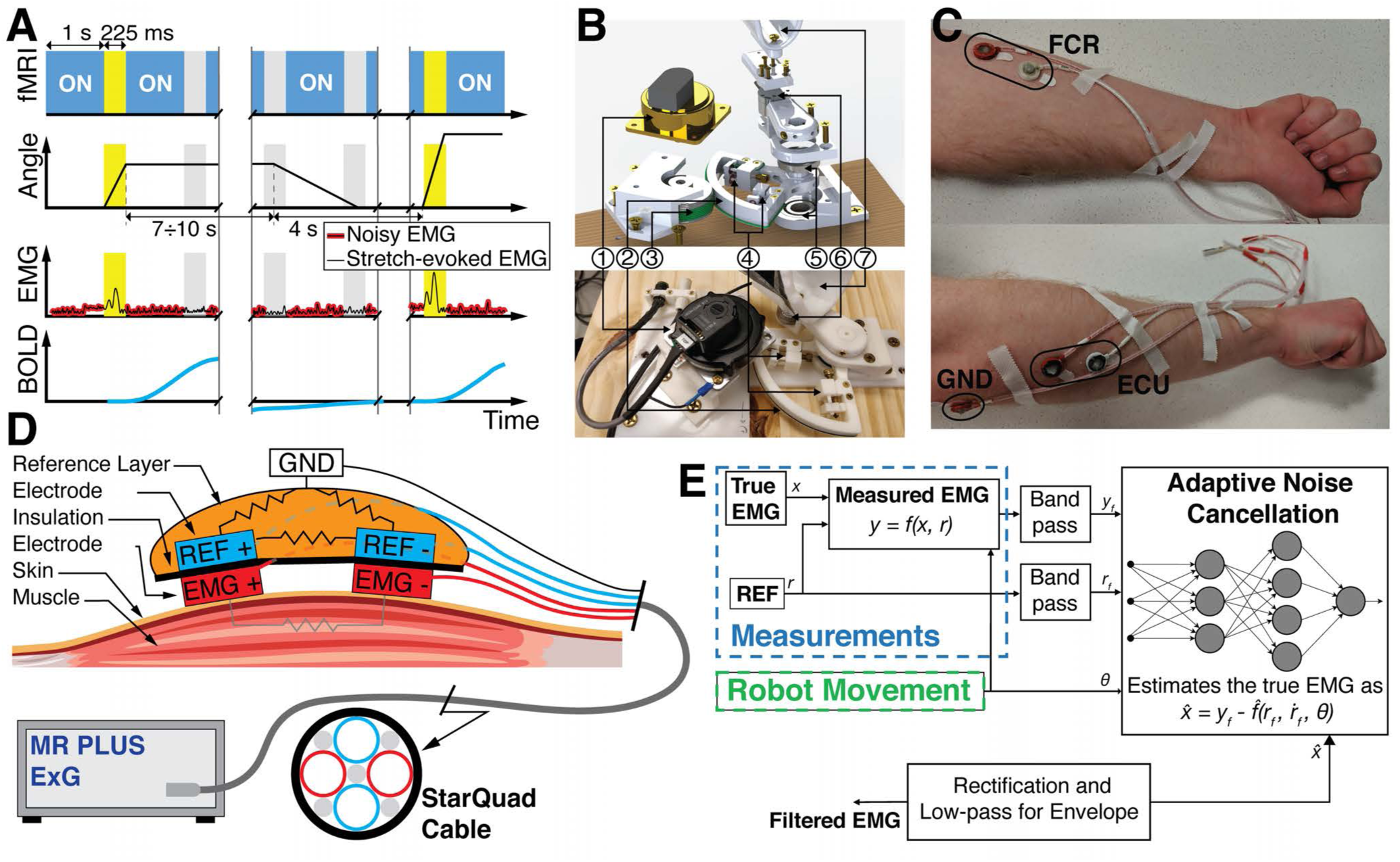
**(A)** Timing diagram of the MRI sequence (top row) with the 225 ms silent windows between acquisition of different volumes; (second row) commanded robot joint trajectory; (third row) expected EMG signal, which is clean during MRI silent windows; (bottom row) expected BOLD signal associated with the LLR; **(B)** Exploded view (top) and prototype (bottom) of the MR-StretchWrist. (1) Ultrasonic motor, (2) output capstan arc, (3) input pulley, (4) tensioning mechanisms, (5) structural bearings, (6) force/torque sensor, (7) hand support; **(C)** Placement of the electrodes on the forearm; **(D)** Hardware of the novel apparatus developed for the acquisition of the EMG data during fMRI protocols **(E)** EMG processing scheme based on Adaptive Noise Cancellation.

Behavioral data from IN and OUT sessions is reported in Sec. Behavioral Results in the supplementary materials. For reference, the mean timeseries of the EMG recorded during Exp 1 and processed using three different filtering pipelines is shown in Fig. 2A.

**Figure 2.**
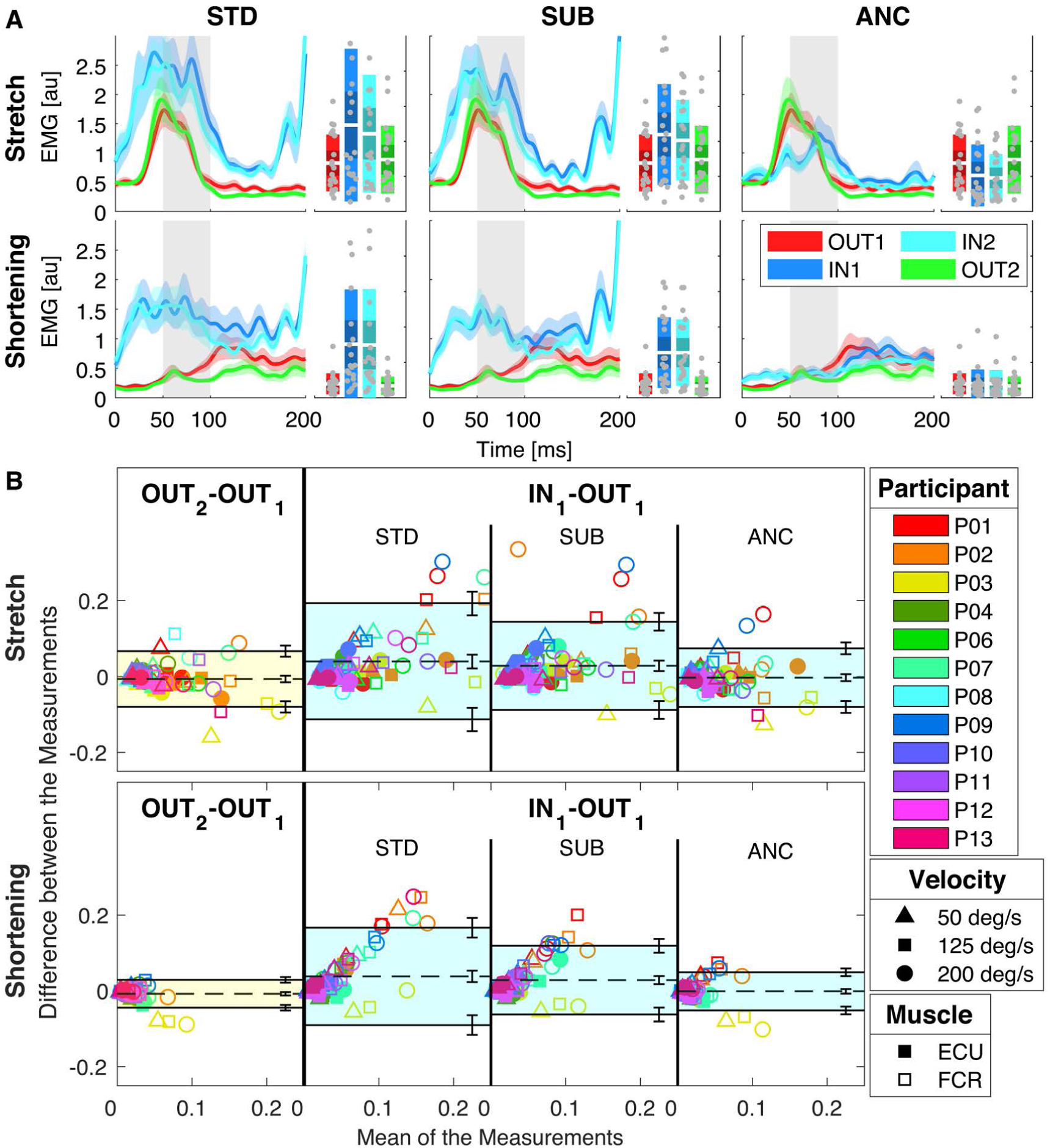
(**A**) Group-average of the stretch-evoked EMG signal measured using different processing pipelines in response to a 200 deg/s ramp-and-hold perturbation for both muscles undergoing stretch or shortening for Experiment 1. Plots are obtained using the same hardware (custom electrode set, see Online Methods) and different processing pipelines: standard (STD), a time-domain subtraction of measured and reference signals (SUB), and the novel adaptive noise cancellation pipeline (ANC). In all graphs, the thick line represents the group mean and the shaded area the 95% confidence interval of the group mean. The gray shaded area indicates the time interval where an LLR is expected. Box plots indicate the distribution of LLR amplitude, with indications for the mean (white horizontal line), mean *±* one st. dev. (dark shaded area), 95% confidence interval (light shaded area), with individual measurements overlaid. (**B**) Bland-Altman plots for the comparisons OUT_2_ vs. OUT_1_ (left) and IN_1_ vs. OUT_1_ when different filtering pipelines are considered (on the right). Datapoints in each of the plots encode participants with colors, perturbation velocities with shapes, and muscles from which the EMG is recorded with fillings. The shaded area represents the interval between the negative and positive limits of agreement, while the dashed line indicates bias. Error bars on the side of each plot represent the 95% confidence intervals by which each parameter is estimated. In both panels, muscle activity is grouped by stimulus direction: muscle stretch on top, and shortening on the bottom. The ANC processing pipeline is clearly superior to the other two methods.

### Reliability of stretch-evoked EMG during fMRI

We validated the stretch-evoked EMG collected during fMRI by quantifying agreement between measurements collected inside and outside the MR scanner (see Online Methods). We used two analyses: one based on the mean LLR responses averaged across multiple repetitions for each set of conditions (group-level analysis), and one based on the analysis of EMG measurements of individual perturbations (perturbation-specific analysis).

#### Group level results

Group-level analysis was performed using the Bland-Altman (BA) method^13,14^, whose plots are shown in Fig. 2B for the comparisons OUT_2_ vs. OUT_1_ and IN_1_ vs. OUT_1_, for different signal filtering pipelines. The values of bias, positive and negative limits of agreement (LoA), with the respective confidence intervals are reported in Tab. S8 in the supplementary materials.

Based on the BA analysis, Adaptive Noise Cancellation (ANC) enables reliable estimation of LLR amplitude during fMRI sessions. In fact, the agreement between LLR amplitudes measured inside and outside the MRI is not worse than the agreement between two OUT sessions. Specifically, the bias estimated for the IN_1_ vs. OUT_1_ comparison was not significantly different from the one estimated for the OUT_2_ vs. OUT_1_ comparison (*z*(72)=0.56, *p* = 0.57 for stretch, *z*(72) = 1.70, *p* = 0.09 for shortening). Additionally, the 95% confidence intervals of the bias estimated for both comparisons, in both muscle stimulus directions, intersect the zero value indicating that the measurement is unbiased (Fig. 3A). Finally, the overlap between the range defined by the LoA for two repeated OUT experiments and the range of LoA for IN vs. OUT experiment, quantified by the Jaccard coefficient, approached perfect overlap (mean ± s.e.m. *J* = 0.843 *±* 0.001 for stretch, *J* = 0.783 *±* 0.001 for shortening).

**Figure 3.**
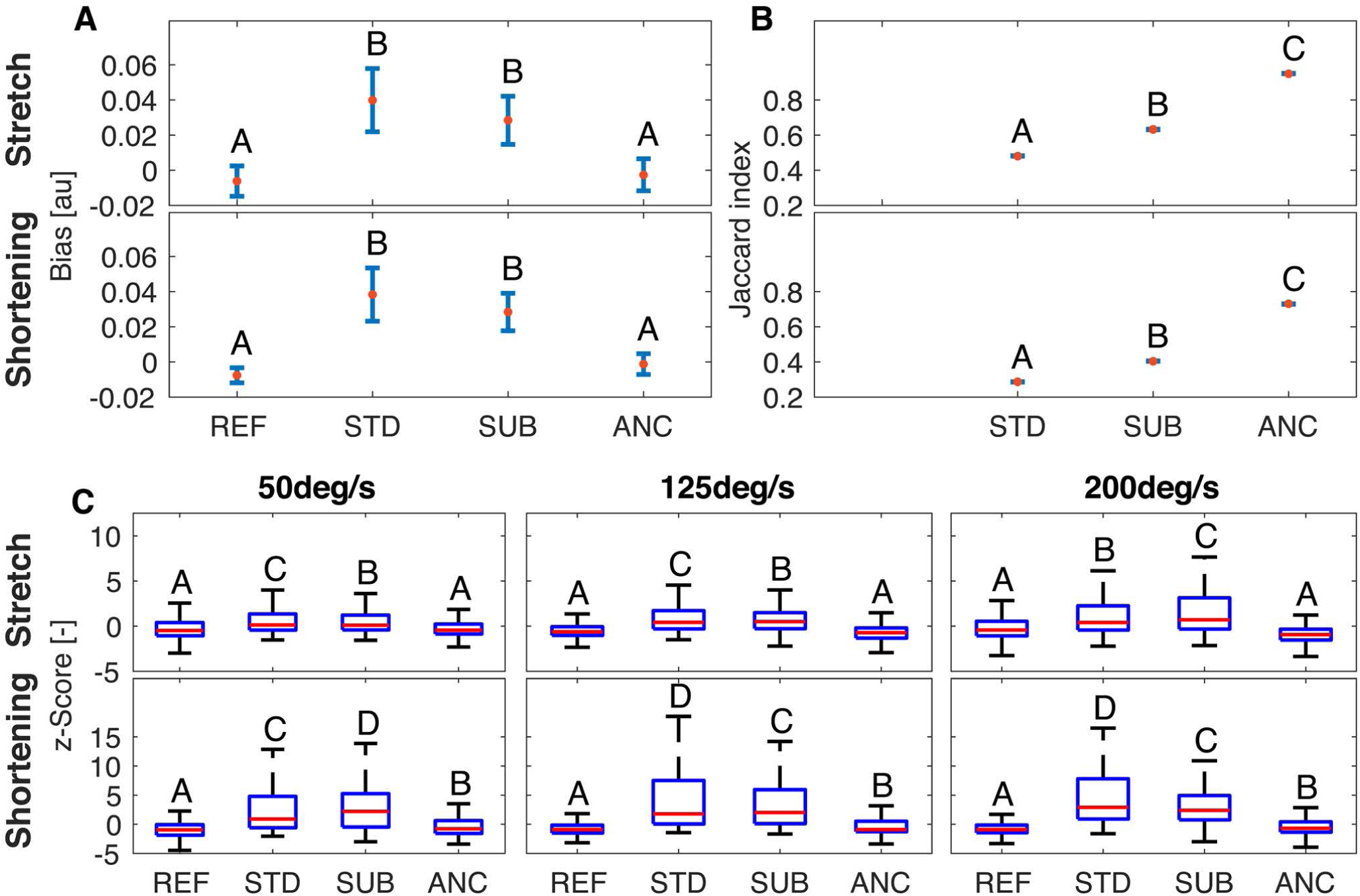
Mean and 95% confidence intervals for the bias (**A**), and the Jaccard index (**B**) obtained for the different filtering pipelines. Test-retest parameters measured for the two stimulus directions are reported in different rows. Letters indicate rankings of the test-retest parameters. Bars that do not share a letter have a statistically different mean. (**C**) Box plots representing the distributions of standardized *z*-scores measured for different combinations of perturbation velocity and filtering pipeline. Distributions obtained for the two stimulus directions are reported in different rows. Letters indicate rankings of the variance of each distribution; boxes that do not share a letter have a statistically different variance.

For reference, test-retest reliability was worse using standard filtering pipelines (STD and SUB - see Online Methods). Specifically, the pairwise comparisons performed between the bias obtained when using different filtering pipelines rejected the null hypothesis for the comparisons STD vs. ANC and SUB vs. ANC, for both stretch (*z*(72) = 3.70, p <0.001; *z*(72) = 4.13, p <0.001, respectively) and shortening (*z*(72) = 4.75, p <0.001; *z*(72) = 4.77, p <0.001, respectively) (Fig. 3A). Bias estimated using ANC was consistently smaller than the one estimated using either STD or SUB (Tab. S8) in both shortening and stretch conditions. The ANC filter also afforded the highest overlap in LoA in quantifying LLRs during fMRI during both muscle stretch and shortening compared to other methods. Jaccard index values for ANC were greater than those obtained with the SUB filter 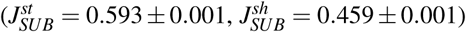, and the STD filter 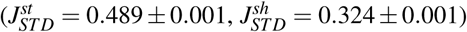. Pairwise comparisons of the Jaccard indices rejected the null hypothesis for all comparisons showing a statistically significant difference between the test-retest error obtained using the different filtering pipelines, in both muscle stimulus conditions (p <0.001 for all comparisons) (Tab. S8, Fig. 3B). The group level analysis has been repeated using measurements collected for the IN_2_ session (Fig. S9, Fig. S10, and Tab. S10, supplementary materials), showing a perfect overlap of all statistically significant results.

#### Perturbation-specific results

To quantify the reliability of measurements of a specific perturbation condition, it is important to measure within-condition variance, something that is not possible using the standard BA analysis. As such, we conducted an additional analysis to quantify the deviation between each stretch-evoked response during IN sessions and the distribution of responses measured in a matched reference (REF) condition (OUT_1_ session), in terms of the *z*-score of each perturbation (Fig. 3C). A Bartlett test established that the variance in measurements collected during IN sessions using ANC was not significantly different from the one during OUT sessions in matched conditions for muscles undergoing stretch (50 deg/s: *χ*^2^ = 3.67, *p* = 0.16; 125 deg/s: *χ*^2^ = 3.69, *p* = 0.16; 200 deg/s: *χ*^2^ = 5.8, *p* = 0.06), while the variance was greater during the IN condition for muscles undergoing shortening (50 deg/s: *χ*^2^ = 17.50, p <0.001; 125 deg/s: *χ*^2^ = 17.87, p <0.001; 200 deg/s: *χ*^2^ = 33.07, p <0.001).

For reference, the variance of *z*-scores measured using other filtering pipelines (STD and SUB) was greater than the one measured for both ANC and REF conditions (*p <* 0.001 for all paired comparisons). Detailed figures and tables arising from this analysis are included in Tab. S9 of the supplementary materials. The perturbation-specific analysis is repeated using measurements collected for the IN_2_ session (Figs. S9-S10, and Tabs. S10-S11 of the supplementary materials), showing a perfect overlap of all statistically significant results.

#### Neural correlates of LLRs

Combining measurements of LLR amplitude measured using EMG with simultaneous whole-brain fMRI, we determined areas of the brain associated with LLRs of specific muscles. The BOLD signal measured from each voxel was analyzed using a general linear model, which partitioned the variance in the measured signal between two regressors of interest and a set of nuisance regressors indicating head movements. Regressors of interests were built as the region-specific hemodynamic impulse response, scaled by the magnitude of LLR amplitude measured for each muscle under stretch (see Online Methods).

We conducted our primary analysis on the brainstem, to test the hypothesis that BOLD signal in any voxel in the brainstem was significantly associated with LLRs for flexors and/or extensor muscles under stretch in the group of individuals tested. We used a small spatial smoothing filter (FWHM = 4 mm) to capture activation of small sets of nuclei, and used a small-volume correction to control for family-wise errors (FWE)^15,16^. Moreover, we conducted an exploratory analysis in the whole brain to further validate the model-based approach pursued, and to analyze function from distributed networks involved in LLRs. For this analysis, we tested whether the BOLD signal in any voxel was significantly associated with LLRs for flexors and/or extensor muscles under stretch. For this exploratory analysis, we used a standard setting for the smoothing filter (FWHM = 8 mm).

To rule out the possibility that measurements of brain activation are not specific to an LLR, but are due to a combination of LLR and background muscle activity applied prior to a perturbation, we conducted an additional analysis based on modeled and measured fMRI data, using a more involved statistical model to describe the BOLD response (see Sec. Supplementary Analyses of the supplementary materials).

#### Brainstem-specific analysis

Muscle-specific activation maps included several clusters of activation in the brainstem. Specifically, LLR-related activity specific to FCR included a total of 57 voxels with supra-threshold *t*-scores, including three distinct clusters spanning bilaterally the midbrain, seven clusters spanning bilaterally the pons, and two clusters in the right superior medulla and inferior pons (Fig. 4 top and Table 1). LLR-related activity specific to ECU included a total of 25 voxels, including three clusters spanning bilaterally the midbrain, two clusters in the left pons and only one voxel in the right pons (Fig. 4 top and Table 1).

**Table 1.**
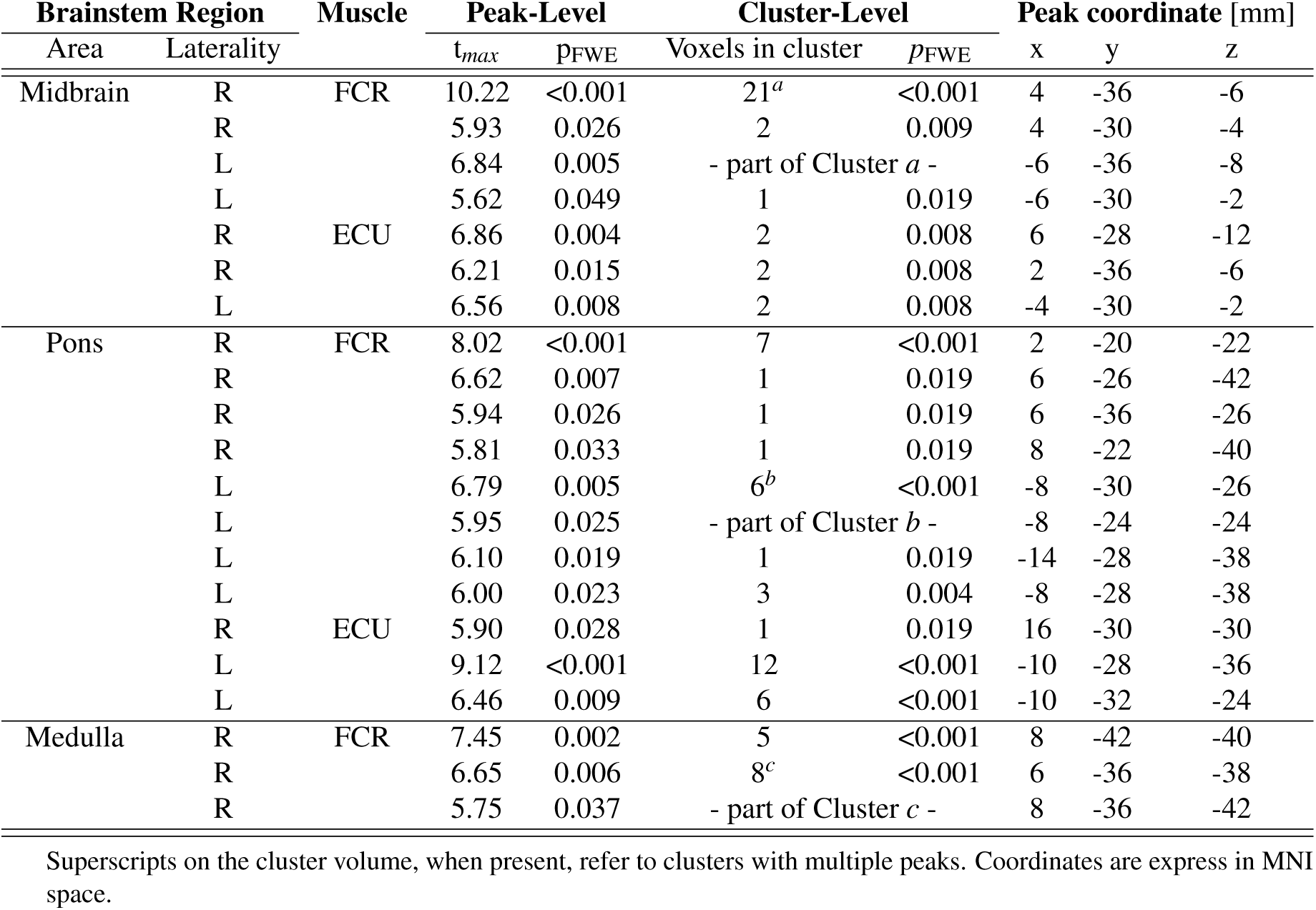
Significance table for the brainstem-specific analysis.

**Figure 4.**
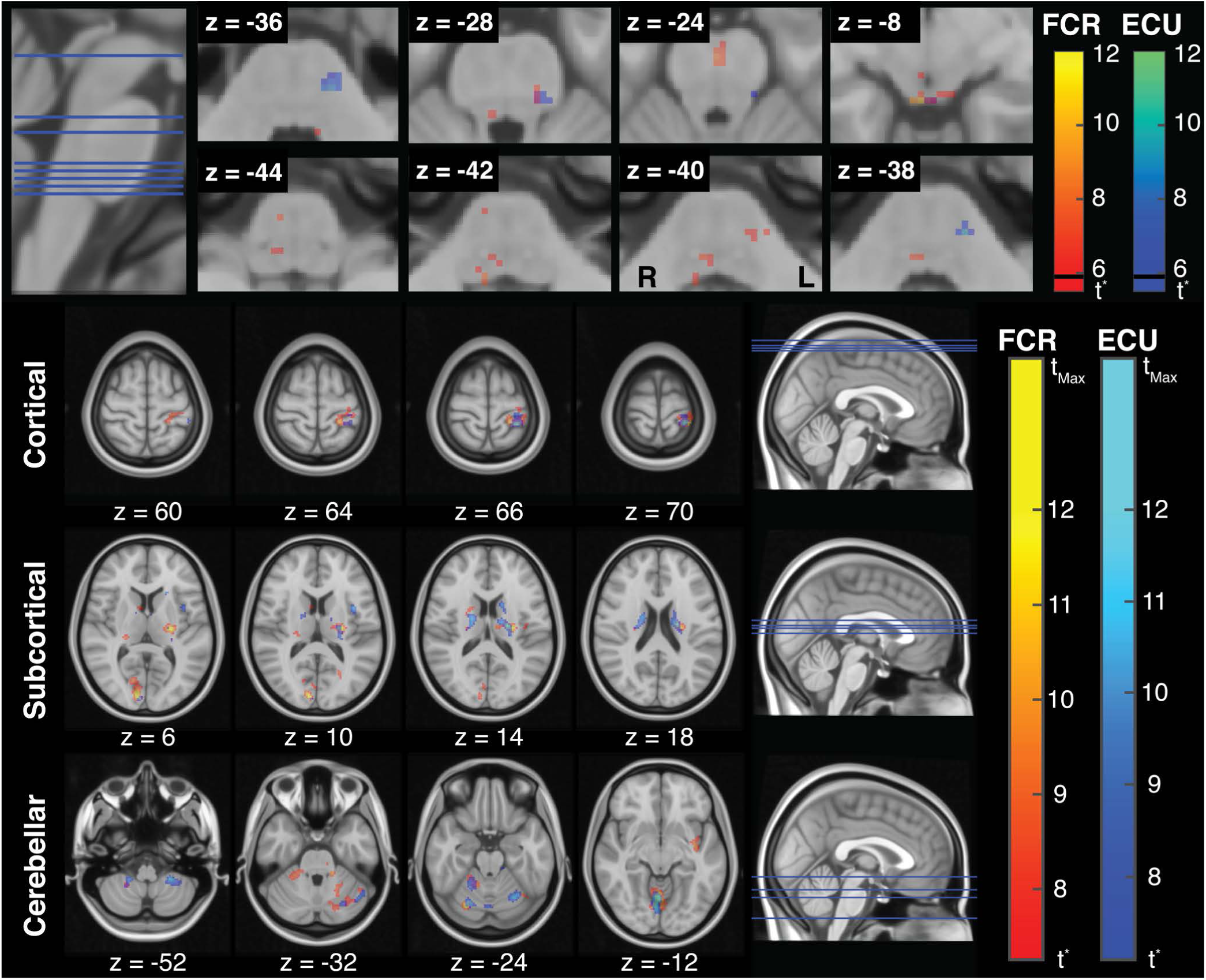
(**Top**) LLR-specific activation maps in the brainstem for FCR and ECU. The maps refer to contrasts *β*_FCR_ > 0 and *β*_ECU_ > 0 obtained for the brainstem-specific analysis. The threshold *t*-statistic, obtained after FWE correction, was equal 5.62 for both regressors. For reference, the t-statistic that refer to a Bonferroni correction (t_Bon_= 5.8) is marked on each colorbar with a black line. (**Bottom**) LLR-specific activation maps in the whole-brain for FCR and ECU. The threshold t-statistic, obtained after FWE correction, was set to 7.25 and 7.09, respectively for FCR and ECU, while t_*Max*_ was equal to 14.95 and 15.54, respectively. Colorbars are saturated at *t* = 12 for better visualization of *t*-statistic gradients. All statistical parametric maps are overlaid on axial slices of the standard Montreal Neurological Institute 152 template, with reported z coordinate in mm

Via subject-specific timeseries analysis of BOLD signal measured from the two clusters with highest statistical scores at the group level, we confirmed the presence of LLR-related BOLD signal specific to FCR stretch in the right medulla and ECU stretch in the left pons (Fig. 5). The subject-specific timeseries analysis suggests that, at least for the clusters of strongest activation at the group level, the dynamics of the measured BOLD signal is more closely associated with LLR dynamics, rather than background activation (see Fig S11 of supplementary materials for more information).

**Figure 5.**
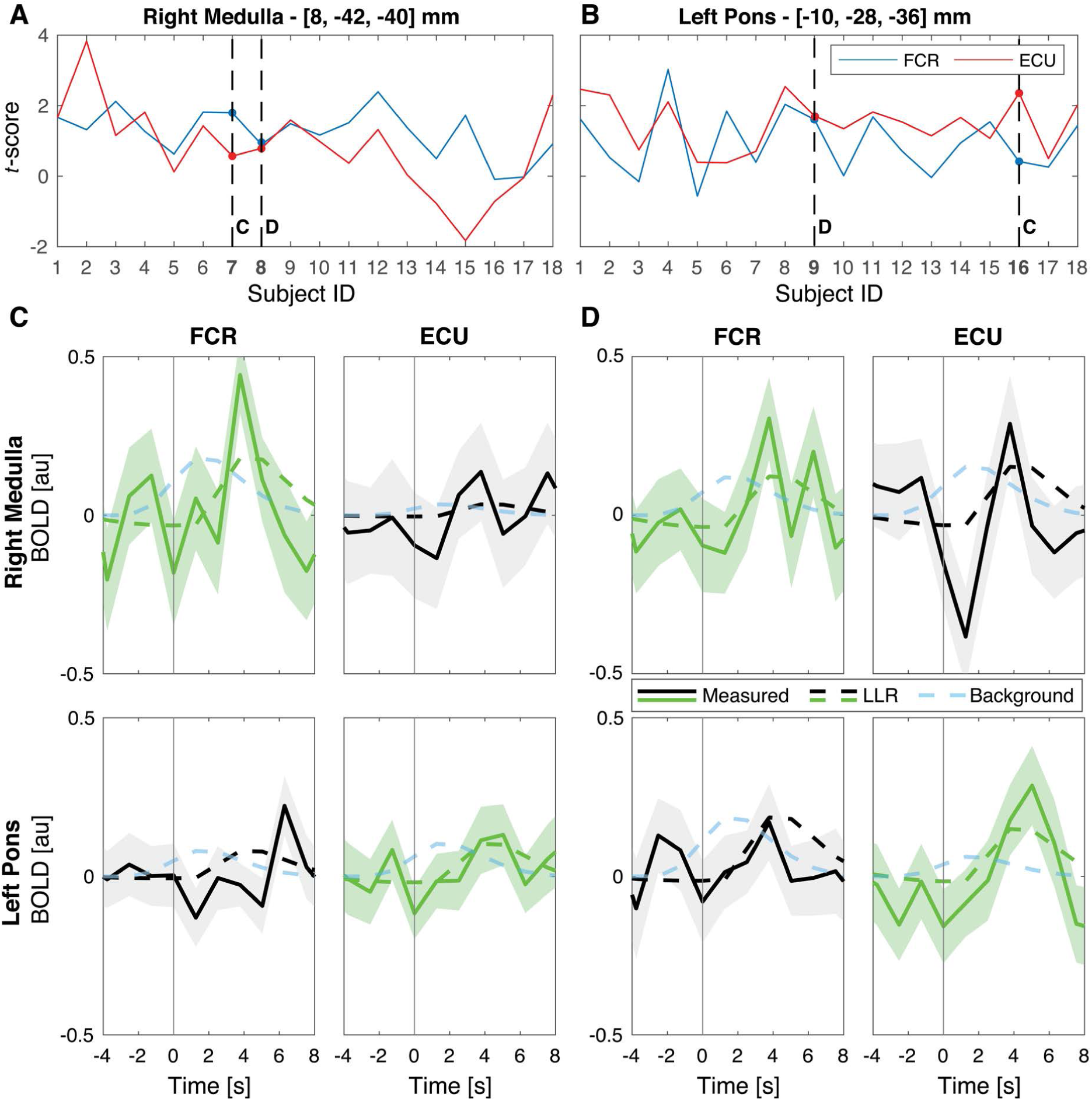
(**A**-**B**) Participant-specific *t* scores obtained for the FCR and ECU activation in the two most significant clusters of activation at the group level in the right medulla (A) and left pons (B). Dashed lines indicate the subjects that have been selected for the timeseries analysis. Specifically, the subjects labeled with **C** showed the greatest difference between the statistical scores specific to the FCR and ECU regressors, while the subjects labeled with **D** showed similar *t* score for the two regressors. (**C**-**D**) Average BOLD signal measured in response to muscle stretch in the selected voxels, for the selected participants. In all graphs, the solid line shows the average residual BOLD response after variance associated to nuisance regressors is removed, while the shaded area represents the 95% confidence interval of the mean response. Dashed lines show the average modeled response respectively for the LLR-specific regressors (green or black line), and for the background-specific regressor (light blue line). Modeled response for the LLR-specific regressor was scaled based on the appropriate *β* coefficient estimated by the GLM model, while modeled background response, which is not included in the primary GLM, was arbitrarily scaled to match the same amplitude of the LLR-specific regressor. Measured and LLR-specific signals are then color-coded to distinguish the response that is significant at the group level for that specific muscle (green), from the one that is not significant (black).

#### Whole-brain analysis

Muscle-specific activation maps included several clusters of activation spanning cortical, subcortical, and cerebellar regions. Specifically, LLR-related cortical activity specific to FCR included contralateral activation in the sensorimotor cortex – specifically, in Brodmann areas (BA) 1, 4a, and 6 –, in the insular cortex (Insula Id1), in the inferior parietal lobule (PFop), and in the opercular cortex (Parietal Operculum OP3), and ipsilateral activation in the intra-parietal Sulcus (hIP2). Additionally, bilateral activation was observed in the inferior parietal lobule (PFcm, PFm on the right hemisphere ans PFop in the left hemisphere), and in the visual cortex (bilateral BA17 and right BA18) (Fig. 4 bottom and Tab. 2). Cerebellar activity included contralateral activation in regions I-IV, VI, and Crus I, and ipsilateral activation in regions V, VI, and VIIIa. Finally, activation could also be observed bilaterally in the thalamus, in the putamen, in the right caudate, and in the right medulla (Fig.4 bottom and Tab. 2).

**Table 2.**
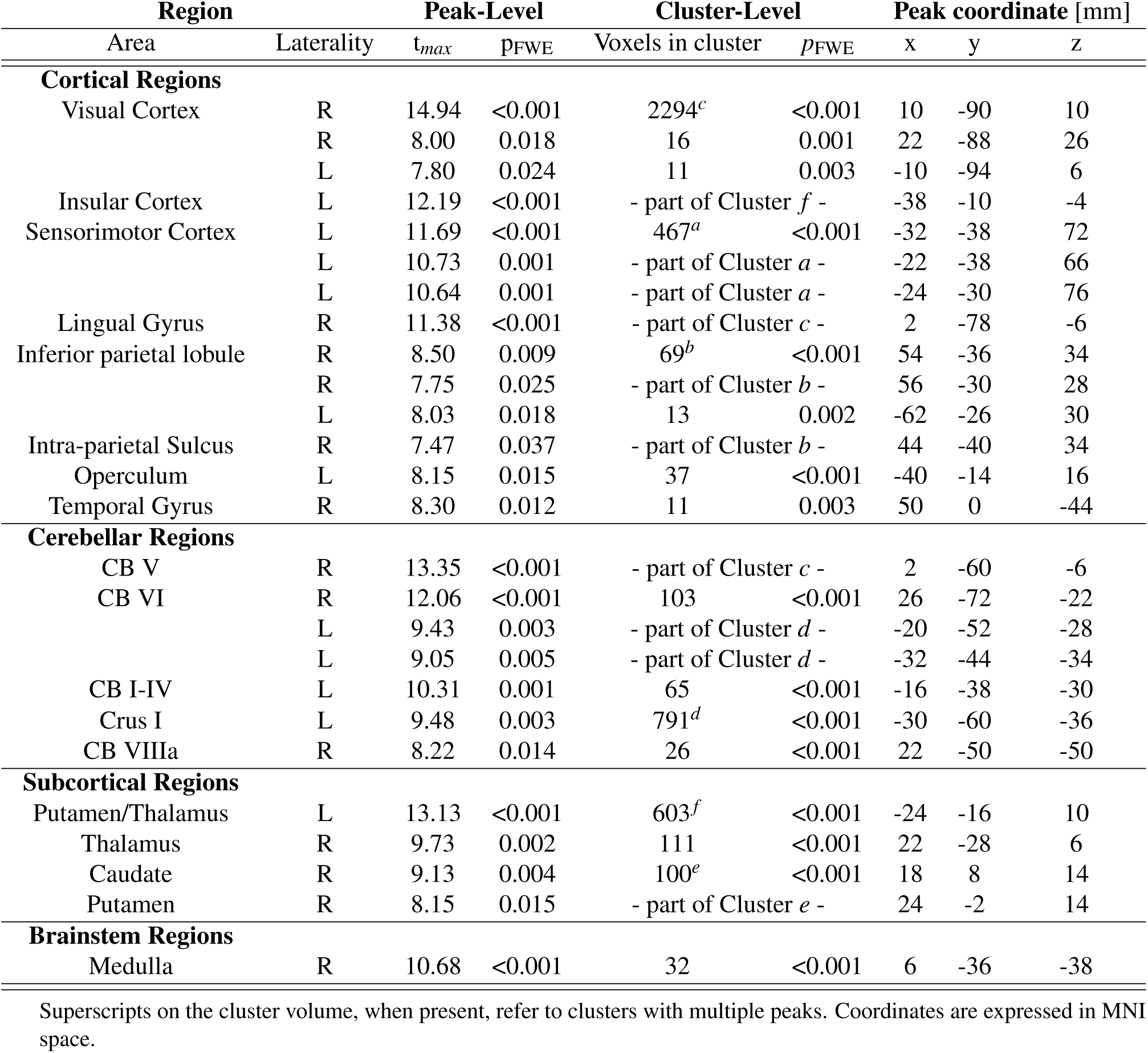
Significance table for the whole-brain analysis for FCR-specific activation.

Similarly, LLR-related cortical activity specific to ECU included contralateral activation in the sensorimotor cortex (BA1, BA4a, and BA6), in the insular cortex (Insula Id1), and in the opercular cortex, and ipsilateral activity in the inferior parietal lobule (PFm), and in the Broca’s Area (BA44) (Fig. 4 bottom and Tab. 3). Cerebellar activity included contralateral activation in regions I-VI, VIII, and Crus I, and ipsilateral activation in regions I-VI, and VIII. Finally, activation could also be observed in the left thalamus and putamen, and in bilaterally in the caudate (Fig. 4 bottom and Tab. 3).

**Table 3.**
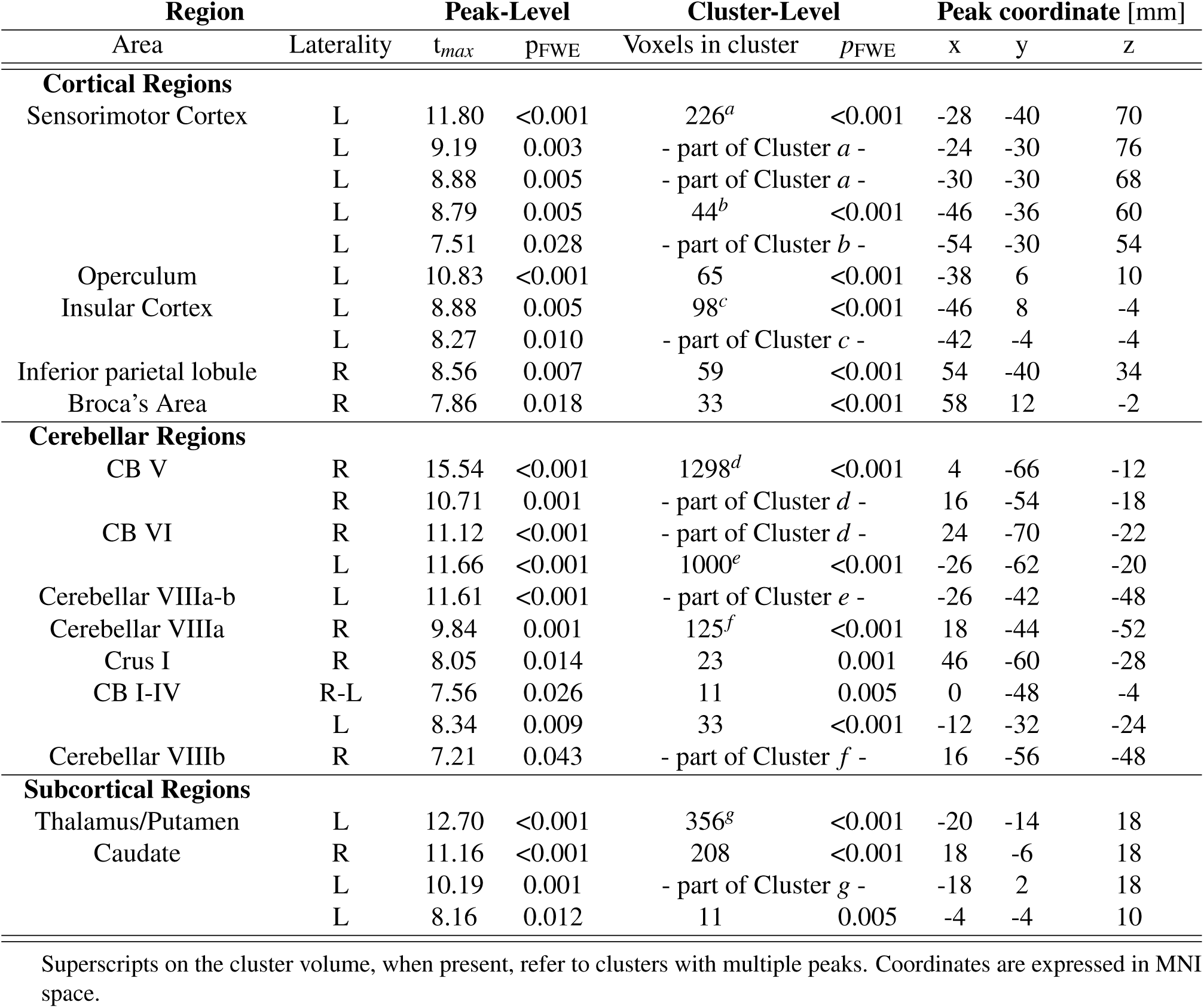
Significance table for the whole-brain analysis for ECU-specific activation.

#### Test-retest reliability of neural activations

By comparing the statistical parametric maps measured in two repeated imaging sessions, we quantified the test-retest reliability of activations in different brain regions (Fig. 6 and Tab. S12). We quantified test-retest reliability both at the group level and at the individual participant level using two metrics, one that quantifies agreement in terms of overlap of thresholded statistical map (Soerensen-Dice index), and one based on intraclass correlation coefficient (ICC).

**Figure 6.**
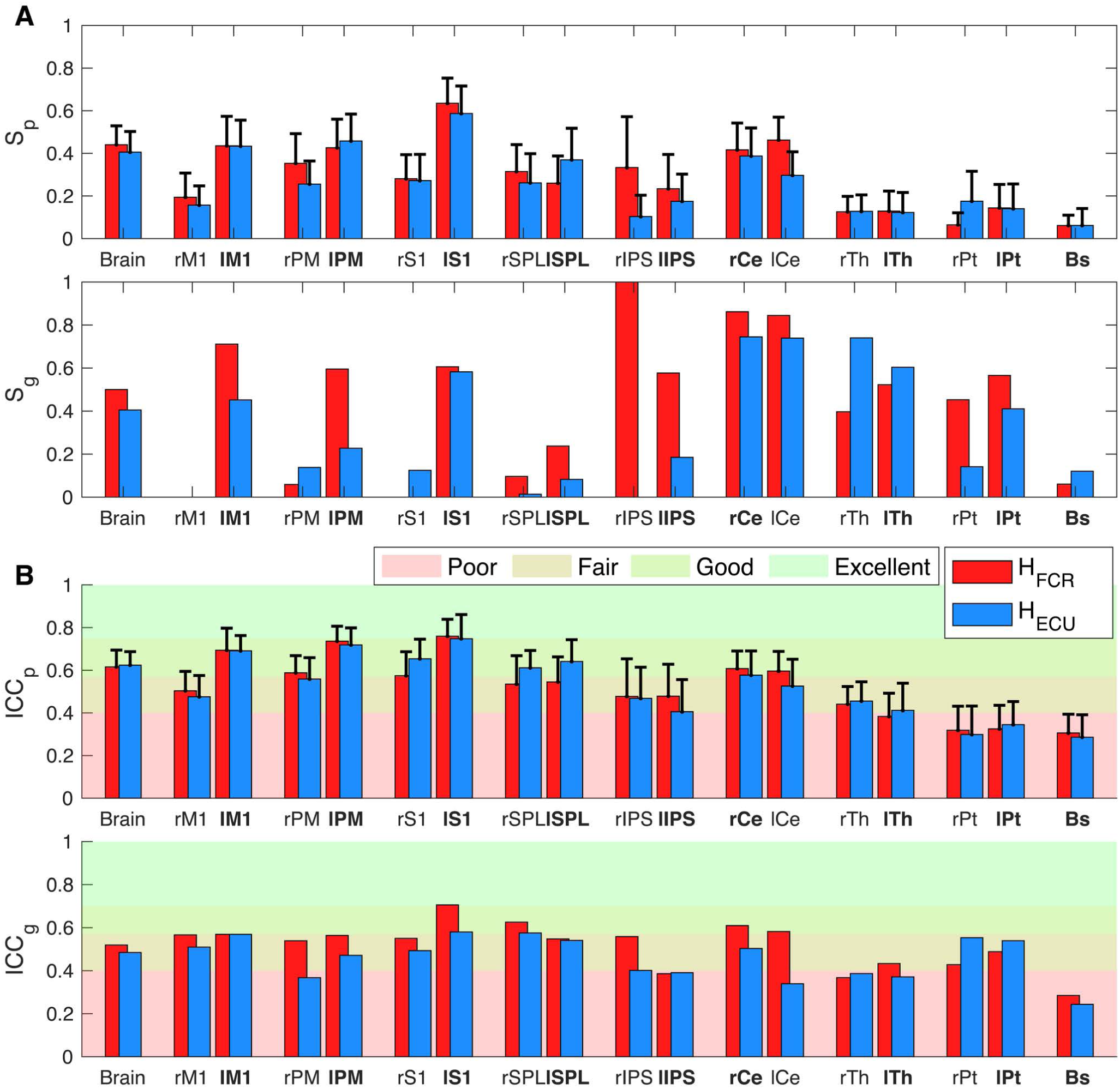
Test-retest reliability scores obtained in the whole brain and in the selected ROIs. (**A**) Sørensen-Dice index, (**B**) Intraclass correlation coefficient. For ICC_*p*_ and S_*p*_, bar height indicates the mean value of the respective metric measured in all participants, with the error bars showing the 95% confidence intervals.

The analysis of the overlap between the group level activation maps showed a moderate to good overlap in the whole brain as well as for all contralateral cortical ROIs for both regressors, with a poor or absent overlap in the ipsilateral ROIs (Fig. 6A and Tab. S12). Good to excellent overlap was observed bilaterally in the cerebellar ROI, while moderate overlap could be observed bilaterally in the Thalamus, in the left Putamen, and in the right Putamen for FCR-specific maps. Poor overlap could instead be observed in the right Putamen for the ECU-specific map and in the brainstem (Fig. 6A and Tab. S12). The analysis of participant-specific activation maps showed a moderate overlap in the whole brain as well as for all selected cortical and cerebellar ROIs, while fair to poor overlap can observed for the subcortical ROIs (Fig. 6A and Tab. S12). Similar reliability was measured for activation associated with flexor and extensor LLRs (paired *t*-test between subject specific Soerensen-Dice index *S*_*p*_ failed to reject the null hypothesis *h*_0_|^1^ that they are the same for all selected selected ROIs with the exception of the right Putamen). Additionally, the paired t-tests between the S_*p*_ measured in the two hemispheres rejected the null hypothesis *h*_0_|^2^ for M1 (FCR: *t*(34) = -2.71, *p* = 0.011; ECU: *t*(34) = -3.44, *p* = 0.003), PM (FCR: *t*(34) = -2.73, *p* = 0.010; ECU: *t*(34) = -2.45, *p* = 0.020), and S1 (FCR: *t*(34) = -2.66, *p* = 0.010), showing a significantly higher overlap in the contralateral sensorimotor cortex for both regressors. No significant difference was observed in any of the cerebellar and subcortical ROIs.

The ICC showed fair to good agreement between the session-specific statistical maps for the cortical and cerebellar ROIs both at the individual subject and the group level (Fig. 6B and Tab. S12). On the contrary, only poor to fair ICC can be observed in the subcortical ROIs (Fig. 6B and Tab. S12). The paired t-tests failed to reject the null hypothesis *h*_0_|^1^ for all ROIs, showing no significantly different test-retest reliability for the two regressors. Similarly to what was observed in the analysis of the overlap of the thresholded maps, the paired t-test rejected the null hypothesis *h*_0_|^2^ for M1 (FCR: *t*(34) = -2.50, *p* = 0.018; ECU: *t*(34) = -3.35, *p* = 0.002), PM (ECU *t*(34) = -2.39, *p* = .022), and S1 (FCR: *t*(30) = -3.99, *p* <0.001; ECU: *t*(34) = -3.41, *p* = 0.002), showing a significantly higher ICC in the contralateral sensorimotor cortex for both regressors. No significant difference was observed in any of the cerebellar and subcortical ROIs.

## Discussion

In this study, we present and validate StretchfMRI, a novel non-invasive technique that enables direct measurement of the subcortical substrates involved in Long-Latency Responses (LLRs) for the first time in humans. StretchfMRI makes a synergistic use of three key components. First, a custom-made MR-compatible wrist robot (the MR-StretchWrist) is used to apply velocity-controlled perturbations at the wrist joint in order to condition LLRs in the wrist muscles. Second, a whole-brain echo-planar fMRI sequence, modified to include a 225 ms silent window after every acquisition volume, is used to obtain gradient artifact-free recordings of the stretch-evoked EMG activity. Finally, a custom EMG electrode set is used in combination with a learning filter to remove movement artifacts from the stretch-evoked EMG activity, thus allowing measurement of reliable EMG data during fMRI scanning. The key new capability afforded by the developed method is the quantification of stretch-evoked muscle responses induced by robotic perturbations using EMG together with fMRI to quantify corresponding neural repsponses. This capability was crucial for establishing the somatotopic arrangement of brainstem activity associated to LLRs of flexors and extensors for the first time non-invasively in humans.

### Quantification of LLR amplitude during fMRI

The use of reference electrodes has been proposed before for processing noisy EEG signals corrupted from noise deriving from either MRI^17^ or motion artifacts induced during walking^18^. However, the sole use of reference electrodes to filter EMG measured during stretch-evoked muscle responses during MRI afforded a test-retest reliability smaller than the one of the physiological process under study (Figs. 2-3, Tabs. S8-S9, and Figs. S9-S10, Tabs. S10-S11). Instead, in our study, we demonstrate that the combination of an MRI-compatible robot to condition LLR responses with Adaptive Noise Cancellation to filter EMG signals with a new electrode set allowed quantification of LLR responses that did not statistically differ from those measured outside the MRI scanner (Figs. 2-3, Tabs. S8-S9, and Figs. S9-S10, Tabs. S10-S11). Moreover, while sEMG has been used during fMRI for isometric tasks^19, 20^ and ocular movements^21, 22^, we demonstrate for the first time reliable sEMG measurements during fMRI for a dynamic task, and quantify their reliability relative to a highly stereotypical physiological process such as Long-Latency Responses.

We combined two analyses to establish the reliability of StretchfMRI in quantifying LLR amplitudes during MRI. We first used a standard method used for test-retest analysis^13, 14^, to quantify bias and limits of agreement of the mean LLR responses averaged across multiple repetitions for each set of conditions, and paired each value with the respective average measurement obtained outside the MRI. Moreover, to isolate the effects of measurement error from those of physiological variability, we defined our outcome measures as contrasts to reference values of reliability obtained when applying the same statistical methods to measurements obtained in two reference sessions, both performed outside the scanner. The results established that our novel filtering pipeline, which uses adaptive noise cancellation to process data recorded during fMRI, grants an agreement with measurements taken outside MRI that is not statistically different from the agreement obtained by repeating the same experiment twice outside MRI (Fig. 3A-B, Tab. S8, and Fig. S10A-B, Tab. S10). Test-retest reliability obtained using alternative processing methods was instead significantly lower than the one obtained for two repeated experiments outside MRI (Fig. 2A-B, Tab. S8, and Fig. S10A-B, Tab. S10).

Because it is important to determine the accuracy in quantifying the LLR for each given perturbation during MRI, we conducted an additional analysis that uses measurements from each specific perturbation. With this analysis, we sought to quantify the deviation between each stretch-evoked response during MRI and the distribution of responses measured in a matched reference condition outside the scanner for each subject. Once again, results demonstrate that only the ANC-based processing pipeline allowed us to achieve, during MRI, a variance of stretch-evoked responses for muscles under stretch (Fig. 3C) that is not different from the one obtained outside MRI (Tab. S9 and Fig. 3C, and Tab. S11 and Fig. S10C).

Overall, these results demonstrate the validity of measurements of LLR amplitude of stretched muscles obtained during fMRI, which allowed us to seek an association between muscle responses and neural function encoded in the BOLD signal.

### Neural correlates of LLRs

The brainstem-specific analysis showed that BOLD signal in the brainstem is modulated by the LLR events in both flexors and extensors. Although previous studies advanced the hypothesis that the brainstem networks should be primarily involved the generation of stretch-evoked responses when task-dependent goals are included in the perturbation protocols^12, 23^, we provide here evidence of stretch-related brainstem activation even in absence of explicit goals (participants were asked to yield to the perturbations). Our results hold true even when using a conservative threshold for quantifying significance of the statistical parametric maps associated with the measured BOLD signal, as the threshold t-statistic obtained with the FWE correction we used is very close to the one resulting from the conservative Bonferroni correction (*t*_*FWE*_ = 5.6 vs. *t*_*Bon*_ = 5.8).

Overall, the muscle-specific statistical parametric maps showed overlapped clusters of activity in the bilateral midbrain, and distinct clusters of activity in the pons and medulla (Fig. 4 and Tab. 1). Specifically, LLR-related activity for FCR included activation in the right superior medulla and bilaterally in the pons. LLR-related activity for ECU included activation only in the left pons. The laterality of observed pontomedullary arrangement is in partial agreement with the currently accepted double reciprocal model of motor functions in the reticulospinal tract^8–10^, with the neural activity of flexors and extensors expected in the right medullary and left pontine reticular formation, respectively. However, the double reciprocal model does not solely describe the complete pattern of activation observed in the brainstem. Clusters of activation observed bilaterally in the pons for FCR are not explained by the model, which suggests that the double reciprocal model is probably not to be intended in an exclusive manner.

Our measurements of brainstem function also include midbrain activity for both FCR and ECU, which is in agreement with previous studies. Taylor et al.^24^ observed how stimulation of the midbrain can modulate the response of the afferent spindle fibers in response to muscle stretch, thus modulating the stretch-reflex threshold^25^. Additionally, the midbrain, together with other subcortical structures such as thalamus, putamen, caudate nucleus, and basal ganglia, is known to play a crucial role in movement initiation^26^, with Cottingham el al.^27^ observing that stimulation of the central gray in the midbrain has a positive effect in facilitating the activation of axial muscles through the reticulospinal pathway.

We performed a supplementary statistical analysis to rule out the possibility that the identified clusters of activation might arise from neural activity associated with background contraction instead of LLR events. The results of this analysis indicate that even after accounting for activation associated with background activity, the LLR-specific activation maps are very similar to the ones obtained in the main analysis. Specifically, statistically significant voxels (Fig. S3 and Tab. S3), while fewer in number, are generally aligned with those identified by the primary GLM in both muscle-specific activation maps (Fig. 4 and Tab. 1). Moreover, the evidence that no supra-threshold voxels associated with the background-specific regressors could be identified in the brainstem, together with the results of the simulation analysis done to quantify specificity and sensitivity of the GLMs, strongly suggest that the clusters of activation included in the brainstem are likely to be associated with true LLR-related activity. Finally, qualitative observation of the timeseries measured in the voxels with highest statistical scores at the group level (Fig. 5) suggest that activation in the right medulla and left pons is indeed primarily associated with LLRs and not with background activity.

The whole-brain exploratory analysis supports the involvement of the cortico-thalamo-cerebellar network for both FCR and ECU. Overlapped clusters of activation can be observed contralaterally in the primary somatosensory cortex, primary motor cortex, and premotor cortex and bilaterally in the thalamus and cerebellum. This observation is in line with our knowledge on motor control^28–30^ and it is in agreement with the findings previously obtained on neural substrates of LLRs^12^. Additional activation could also be observed in secondary somatosensory areas–i.e. left parietal operculum, left insular cortex– for both FCR and ECU. Both areas are part of the somatosensory system and have been observed to be involved in processing light touch sensation, tactile attention^31, 32^, and the sense of bodily-ownership and proprioception^33–35^. Finally, overlapped clusters of activation referring to the two muscles can be observed in the right caudate nucleus, left putamen, and the right inferior parietal lobule. While the involvement of the inferior parietal lobule in controlling motor actions is still under debate, with few studies that propose its involvement in coding different motor actions^36, 37^, the role of putamen and caudate nuclei is much more established^26, 38, 39^. Both subcortical regions have been observed to be involved in postural maintenance and sensory-motor integration^40^ and in controlling movement initiation^26^.

fMRI activation data obtained using StretchfMRI afford test-retest reliability that varies according to different brain regions. In terms of a standard reliability scale^41^ used in several other fMRI test-retest reliability studies^42–46^, we observed moderate to good reliability in the whole brain as well as for all cortical and cerebellar ROIs both at the subject and the group level (Tab. S12 and Fig. 6). On the other hand, we observed lower reliability in subcortical ROIs, with poor overlap and poor to fair ICC (Tab. S12 and Fig. 6). While the reliability for the brainstem is low, it is important to consider two main factors. First, event-related designs, which are necessary to associate stretch-evoked muscle activity with BOLD signal, are known to have a lower test-retest reliability compared to a standard block design^47^. Second, the smaller FWHM of the smoothing filter used for the brainstem-specific analysis^48, 49^ negatively affected the test-retest reliability metrics^42^. As a comparison, the ICC_*g*_ score quantified for the brainstem changed from 0.28 to 0.43 and 0.24 to 0.39 for FCR and ECU, respectively. As such, low test-retest reliability presented here is obtained in light of increasing sensitivity to capture small regional variations of nuclei function. To the best of our knowledge, no previous methods studied function in the brainstem using event-related protocols, nor reported values of test-retest reliability for such methods. Nonetheless, in all ROIs the reliability metrics assume values that are comparable with those reported by previous studies for both cortical^42–44, 50^ and subcortical ROIs^45, 46^.

### Conclusions

In conclusion, in this work we presented and validated StretchfMRI, a novel technique that for the first time enables *in-vivo* measurements of brainstem function during stretch-evoked responses in humans. Statistical parametric maps of muscle-specific responses in the brainstem in part support established models of the organization of motor function in the pontomedullary reticular formation^8–10^, showing how excitatory stretch-evoked responses in the flexors and extensors correlate with neural activity in the ipsilateral medulla and contralateral pons, respectively. StretchfMRI allows for the first time quantitative analysis of function in a secondary motor pathway, the reticulospinal tract (RST), thus enabling new investigations in both basic and translational neuroscience. As an example, StretchfMRI can be used to further investigate the task-dependent modulation of brainstem activity associated with LLRs proposed in^51–53^. Because function in the RST has been advanced to play a role in post-stroke recovery or impairment^54, 55^, but measured indirectly and with limited spatial specificity, it is possible that StretchfMRI will result in an improved understanding of mechanisms of recovery and impairment from CST lesions.

## Acknowledgments

Research reported in this publication was supported by the National Institute Of Neurological Disorders And Stroke of the National Institutes of Health under Award Number R21NS111310. The content is solely the responsibility of the authors and does not necessarily represent the official views of the National Institutes of Health. Additional support was provided by the University of Delaware Research Foundation grant no. 16A01402, from ACCEL NIGMS IDeA grant no. U54-GM104941.

## Author contributions statement

A.Z. and F.S. developed StretchfMRI, designed experiments and data analyses. A.Z. conducted the experiments. A.J.F. and D.R. supported neuroimaging data analysis. A.Z. implemented all data analyses. All authors reviewed the manuscript.

## Additional information

To include, in this order: **Accession codes** (where applicable);

**Competing interests** (mandatory statement).

The corresponding author is responsible for submitting a competing interests statement on behalf of all authors of the paper. This statement must be included in the submitted article file.

## Online methods

### StretchfMRI Technique

The investigation of the neural substrates of LLRs via fMRI requires simultaneous measurement of muscular and neural activity during the application of velocity-controlled perturbations that condition LLRs in a set of muscles. While previous studies primarily investigated reflex activity of proximal muscles applying perturbations to the shoulder and elbow joints^12^, we focused our attention to forearm muscles during wrist perturbations. By targeting wrist movements, we expect to have reduced head and body motion, two factors that can degrade the quality of the fMRI images^56^ and of EMG recordings^19, 57^.

### MR-StretchWrist device

Previous studies observed that LLRs can be elicited in the forearm muscles with background wrist torque and perturbation velocities that range from 0 to 0.5 Nm and from 100 to 250 deg/s, respectively^52, 58–61^. To determine the gearmotor torque characteristic required to elicit long-latency responses, we modeled the wrist joint as a mass-spring system^62^, assuming the hand inertia to be *I*_*h*_ = 0.0024 kg m^62^ and considering the stiffness resulting from the short-range response of muscle fibers (short-range stiffness model used in our previous work^63^), and requiring the hand to reach a target velocity of 250 deg/s within 50 ms (time considered as the onset of LLRs). With these assumptions, we determined that a peak torque *τ*_*R*_ = 3 Nm would be required when a constant background torque of 0.5 Nm is applied.

To apply such perturbations, we have developed a novel robotic device, the MR-StretchWrist (MR-SW) shown in Fig. 1B. The MR-SW is a 1-Degree of Freedom (DOF) robot capable of applying controlled rotations of the hand about the the wrist Flexion/Extension (FE) axis in a range of *θ*_*FE*_ = [−45; 45] deg. It is actuated by an ultrasonic piezoelectric motor (EN60 motor, Shinsei Motor Inc., Japan) which provides 1 Nm peak torque and 900 deg/s peak velocity, previously utilized for several applications in MR-compatible robotics^64–66^. To fulfill the design specifications, we have employed a capstan transmission with 3:1 gear ratio to transfer motion from the motor to the end effector. The capstan drive consists of two pulleys with different diameters connected by a smooth cable that is wrapped around the pulleys multiple times to ensure no-slippage high-friction contact (Fig. 1B). Such transmission is ideal for this application as it is characterized by no backlash, low friction, and high bandwidth and is easily manufacturable using MR-compatible materials. The MR-SW is instrumented with a six-axis MR-compatible Force/Toque sensor (Mini27Ti, ATI Industrial Automation, Apex, NC) to measure the wrist joint torque.

To ensure MR-compatibility of the entire system, all structural components have been manufactured using ABS and connected using brass screws. For the cable of the capstan transmission, we have used a microfiber braided line (SpiderWire Stealth SPW-0039, 0.4 mm diameter braided fishing line). The output shaft is supported by ceramic radial bearings (Boca Bearings, Boynton Beach, FL, USA), while brass shafts are included to support the tensioning mechanisms. To reduce electromagnetic interference introduced in the MRI scanner room by the motor and the motor encoder, a tripolar twisted-pair shielded cable was used for encoder line and shielded cable was used for the motor power line. Both lines are filtered when passing through the scanner patch panel using 5.6 pF and 1.3 pF capacitive filters, respectively. We have assessed the MR-compatibility of the MR-SW in a previous study^67^ following the methods described in^68^.

### Simultaneous recording of fMRI and EMG data

Measuring EMG during fMRI protocols is technically challenging because of the large artifacts introduced in the EMG recordings by the coupling of the fMRI time- and spatially-varying electromagnetic fields–i.e. static magnetic field, gradient magnetic field, radio wave–with the undesired movement of the EMG electrodes. While the radio waves introduce noise at a frequency range that is distinct from the spectrum expected for physiological muscle contractions, meaning that noise could be removed using frequency domain filters, this is not the case for gradient and movement artifacts whose spectrum overlaps with the one expected for muscle contractions.

Different methods have been proposed to compensate for the gradient artifacts in electroencephalogram (EEG) and EMG protocols^57, 69–71^ collected during MRI. These methods are mainly based on template removal algorithms that subtract from the EEG/EMG recording the average signal recorded across several volumes to estimate an artifact-free biosignal. Template subtraction is based on the assumption that the artifact waveforms repeat equally for all collected volumes. However, this assumption fails when artifacts are coupled with highly variable movement of the electrodes. Given our interest in measuring EMG only during brief periods of time–i.e. those where we expect to observe a LLR–we pursued a simplified approach to avoid gradient artifacts. We introduced in the MRI scanning protocol a 225 ms silent window after each acquisition volume where no fMRI excitation is generated. By synchronizing the application of the velocity-controlled perturbations with the timing of the fMRI sequence, it was then possible to obtain gradient artifact-free measurements of LLR (Fig. 1A). Doing so, we leveraged the intrinsic delay of the hemodynamic signal that decouples temporally the measurement of muscle activity associated with an LLR–expected within 100 ms of a perturbation–with the measurement of the associated blood-oxygen level dependent (BOLD) signal–which completes its course several seconds after a reflex is elicited (Fig. 1A).

While this approach allowed us to remove artifacts associated with both radio waves and gradients, preliminary analyses have highlighted that sub-millimeter movements of the electrodes caused by the robotic perturbation, while tolerable in normal laboratory conditions, produce artifacts up to 5-10 times larger than the stretch-evoked EMG response when they are coupled with the magnet’s static magnetic field (Fig. S5 in the Supplementary Materials).

These artifacts are a consequence of the Maxwell-Faraday law that describes how a time-varying magnetic flux through the surface enclosed by a conductive loop induces current on the conductive loop. For our specific application, the time-varying flux is generated mainly as a consequence of the fact that the path of the electrode leads during wrist perturbation unpredictably moves and deforms, leading to a change in the area enclosed by the conductive loop. Moreover, because the MRI static magnetic field is non-homogeneous in space, the amplitude of the artifact is highly dependent on the position that the electrodes occupy in space, which cannot be guaranteed to be constant during an experiment.

However, since overall motion-induced artifacts are ultimately a function of the position of electrodes and wire path over time, they could theoretically be removed by 1) co-locating reference electrodes (measuring only motion-induced artifacts) with measurement electrodes (measuring both motion-induced artifacts and muscle EMG), 2) matching the electrode-to-electrode and electrode-to-ground impedance of the two sets of electrodes, 3) routing the leads of the reference and measurement electrodes so that their path is the same.

We have developed a novel electrode set that embodies these principles to measure EMG signal of forearm muscles during fMRI (Fig. 1D). The set includes a bipolar reference electrode (REF) separated by a measurement electrode (EMG) via an insulating layer. Both couple of electrodes are off-the-shelf Ag/AgCl bipolar electrodes (multitrode, Brain Products, Munich, Germany), with the terminals of the EMG electrode that are placed on the belly of the corresponding muscle and aligned with the direction of the muscle fiber. The REF electrode pair, colocated with the EMG pair, is then embedded in a conductive substrate (Squishy Circuits, Anoka, MN, USA) that approximates the electrode-to-electrode and electrode-to-ground impedance of superficial forearm muscles (2-5 MΩ). To obtain the best matching wire path between REF and EMG electrode leads, all wires are routed through a StarQuad cable (VDC 268-026-000, Van Damme Cable). Proper coiling around a central core in a StarQuad cable ensures that differential signals of the two sets of bipolar electrodes carry the same motion-induced signal of a virtual conductor routed through the center of the core. The conductive substrate was then grounded using the shield terminal of the StarQuad cable to the amplifier ground, and connected to a fifth Ag/AgCl electrode placed on the lateral epicondyle of the elbow. A similar approach has been used by^17^ to minimize fMRI-induced artifacts on EEG recordings.

Because perfect (i.e. whole spectrum) impedance matching cannot be guaranteed, a simple subtraction of the EMG and REF signals would not ensure sufficient compensation of motion artifacts. To solve this issue, we developed a novel processing scheme that uses adaptive noise cancellation (ANC) to estimate artifact-free EMG signal (Fig. 1F). ANC is an adaptive filtering technique based on a multi-layer neural network that can be used to estimate a desired signal *x* corrupted by an interference *w*^72–75^. To properly work, the ANC takes two inputs consisting of the corrupted signal *y* = *x* + *w* and a measurement of one or more interference signals **r**. Considering that the true interference is altered when passing through two different measurement channels, signals **r** and *w* may be non-linearly related, such that a simple subtraction between *y* and components of **r** usually causes the distortion of the estimated signal 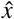. To solve this problem, ANC attempts to learn the non-linear relationship *f* (**r**) between the interference measured by the two different channels, estimating the signal *ŵ*= *f* (**r**), so that the desired signal can be estimated as 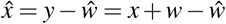.

For our analysis, we implemented the ANC scheme in MATLAB 2019a (MathWorks Inc., Natick, MA, USA) using an Artificial Neural Network Fuzzy Inference System (ANFIS)^76^. After appropriate learning, ANFIS constructs the non-linear mapping for the input-output relationship (estimate *x* given **r** and *y* in our case) without requiring apriori knowledge of the structure of *f* (**r**). Learning occurs by tuning a set of membership functions that refer to the different layers of the network. In our algorithm, we used a hybrid iterative optimization method consisting of back-propagation for the parameters associated with the input membership functions, and least squares estimation for the parameters associated with the output membership functions^76^. Since the model structure used for the model has a large number of parameters, there is a risk for the ANFIS to overfit the data. To avoid overfitting, the algorithm partitions at every iteration a random sample from the entire dataset (in our case, a single continuous random interval within the time-series of *y* and **r** collected during a given perturbation) and uses it to cross-validate the model. The idea is that the cross validation error decreases until overfitting starts to occur. As such, the algorithm selects the set of parameters for the membership functions that refer to the solution that is characterized with the minimum cross-validation error^76^.

In our implementation, we considered the interference signals **r** as a three-component vector, including the interference signal *r* measured by REF electrodes, its derivative 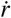, and a signal that quantifies the perturbation-related motion, as measured by the rotative encoder built in the MR-SW (*θ*). The ANFIS was implemented using the *genfis* and *anfis* functions built in the Fuzzy Logic Toolbox.

### Experimental Methods

#### Population

27 healthy individuals (16 males, 11 females; age range 19-38 years) volunteered to participate in one of the two experimental protocols that compose this study. 13 participants were involved in Experiment 1, 14 participants were involved in Experiment 2. All participants self reported as right handed, free from neurological disorders, orthopedic or pain conditions affecting the right arm and provided informed written consent prior to data collection. This study was approved by the Investigation Review Board of the University of Delaware Protocol no. 1097082-5 and was conducted in accordance with the Declaration of Helsinki.

#### Experimental procedures

In both protocols, participants were exposed to a sequence of Ramp-and-Hold (RaH) perturbations in either flexion or extension at multiple velocities while EMG was recorded from the flexor carpi radialis (FCR) and the extensor carpi ulnaris (ECU) (Fig. 1C), as described in Sec. Simultaneous recording of fMRI and EMG data. Before the perturbation onset, participants were visually cued to apply 200 mNm of background torque in the direction opposing the ensuing perturbation (e.g. if the perturbation was in the extension direction the participant was required to apply a flexion torque) to condition the muscles that would be stretched by the perturbation. The perturbation was then automatically triggered after the error between the target and measured torque was below 25 mNm for a given time T_hold_. To avoid habituation to the time delay, for each perturbation the time T_hold_ was randomly selected between 400 ms and 800 ms. Because the impact dynamics that characterize the end of a perturbation generates undesired oscillations in the EMG recording, leading to detection of false positive activity in the LLR window, perturbations were kept active for a fixed time of 200 ms, regardless of the magnitude of perturbation velocity. A rest period lasting for an interval randomly selected between 7 s and 10 s was then included after each perturbation^77^, after which the robot slowly moved the hand back to the neutral position with a return velocity set to 25 deg/s. Additional 4 s of rest were then allowed before the toque target for the following perturbation was displayed to the participant. Participants were instructed to relax their muscle and yield to all perturbations throughout the entire experimental protocol.

In each session, participants were exposed to a total of 60 perturbations, pseudo-randomly selected from a pool of six different perturbation velocities ([50 125 200] deg/s in flexion or extension) repeated 10 times each. In both experiments, before the beginning of each session, participants were visually cued to apply and hold a set of ten interleaved flexion and extension isometric toques. In both directions, the magnitude of the desired torque was set to be 500 mNm, and the participants were asked to keep it constant with a maximum error of 25 mNm for 5 s. This set of contractions was used to normalize stretch-evoked responses and enable group analysis of collected EMG data.

All experiments were conducted at the Center for Biomedical and Brain Imaging located at University of Delaware using a 64 channel head coil on a Siemens Prisma 3T scanner.

##### Experiment 1

Experiment 1 was composed of a total of four sessions, all performed during the same visit. The first and last sessions (OUT_1_ and OUT_2_) were performed in a mock scanner outside the MRI room, allowing the participant to maintain a supine posture similar to the one required for fMRI scanning. This was done in the attempt to replicate imaging conditions as much as possible, while removing any source of interference induced by the MRI electromagnetic fields. The two middle sessions (IN_1_ and IN_2_) were performed inside the MRI scanner during fMRI scanning using the protocol described in Sec. Simultaneous recording of fMRI and EMG data. Parameters used for the MRI sequence included: Multi-Band Accelerated EPI Pulse sequence; 2×2×2 mm^3^ voxel resolution with 0.3 mm slice spacing, 46 deg degree flip angle, 880×880 px per image, 64×110×110 mm^3^ image volume; TR=1225 ms, and TE=30 ms; pixel bandwidth=1625 Hz/pixel, receiver gain: high. After the two functional sessions, a high resolution structural scan (magnetization-prepared rapid acquisition with gradient echo (MPRAGE): 0.7×0.7×0.7 mm^3^ resolution, with TR=2300 ms, and TE=3.2 ms; 160 slices with a 256×256 px per image, for a 160×256×256 mm^3^ image volume) was acquired for registration and normalization of the results to a common space. The full experimental protocol took about 140 minutes divided as 45 minutes for the robot setup and the electrode placement, 20 minutes per each one of the four experimental sessions, and 15 minutes for the structural scan.

##### Experiment 2

Experiment 2 was composed by only sessions IN_1_ and IN_2_ performed in Experiment 1–i.e. those performed during fMRI scanning. The full experimental protocol took about 85 minutes divided as 30 minutes for the robot setup and the electrode placement, 20 minutes per each one of the two experimental sessions, and 15 minutes for the structural scan.

#### EMG acquisition and analysis

Surface EMG was recorded with the BrainVision Recorder software (Brain Products, Munich, Germany) using a 16-channel MR-compatible bipolar amplifier (ExG, Brain Products, Munich, Germany). Given our interest in studying the neural substrates of LLR for both flexors and extensors, EMG was recorded from two representative wrist muscles: Flexor Carpi Radialis (FCR) and Extensor Carpi Ulnaris (ECU) (Fig. 1C).

For each muscle, after having carefully cleaned the skin with a 70% Isopropyl Alcohol solution, we placed the bottom layer of electrodes on the belly of the muscle oriented along the muscle fiber, and filled the central hole of each electrode with an abrasive Electrolyte-Gel (Abralyt HiCl, Rouge Resolution, Cardiff, UK). Contact impedance for each electrode was measured using the BrainVision Recoder software and a cotton swab dipped in abrasive gel was swirled on the skin until the measured contact impedance was lower then 10 kΩ, as described in the product technical specification. We then carefully co-located the reference electrodes on top of the measurement electrodes, using a layer of electric tape applied on top of the measurement electrodes to avoid electrical contact. Finally, we placed the conductive substrate on top of the reference electrodes and connected it to ground. In order to minimize relative motion of the different components of the apparatus, we applied pre-wrap around the entire forearm.

EMG signal was processed using a standard pipeline^51, >78^, modified to include the ANC or the SUB pipelines. Both REF and EMG signals were initially segmented to extract the subset of data points representing perturbation-related activity recorded during the 200 ms silent window (25 ms after volume acquisition is completed), so that the first time point corresponds to the perturbation onset. The segmented signals were then band-pass filtered using a 4^th^ order Butterworth filter with cut-off frequencies *f*_*LP*_ = 20 Hz, and *f*_*HP*_ = 250 Hz, and fed to the later components of the filtering pipeline (as described in Fig. 1E). The estimate of the EMG activity returned by the ANC, SUB, and STD filters was finally rectified and low-pass filtered with a 4^th^ order Butterworth filter with cut-off frequency *f*_*ENV*_ = 60 Hz. To allow between-subject and between-muscle comparison, after filtering, we normalized the stretch-evoked EMG activity by the average EMG 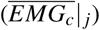 measured during the isometric contractions of the muscle *j* recorded prior to the beginning of each perturbation session. To determine 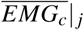, we used only the central 3 s of activity recorded for the subset of contractions in which the given muscle was active–i.e. only the flexion torques for the FCR and only the extension torques for the ECU. The same constant was used to normalize EMG activity measured in response to perturbations that both stretch and shorten the muscle. Finally, to extract the magnitude of the long-latency response *H*_*i, j*_ elicited by perturbation *i* on muscle *j*, we used the *cumsum* method^79^, quantifying *H*_*i, j*_ as the area underlying the processed EMG signal *EMG*_*i, j*_(*t*) in the temporal window [50, 100] ms after the perturbation onset:

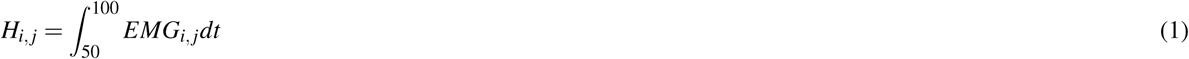

where time is expressed in ms.

#### Validation of the EMG measurements during fMRI

To validate the ability of StretchfMRI to reliably condition and quantify stretch-evoked LLR responses during fMRI, we used the EMG data recorded in the four sessions of Experiment 1. As no MR-related noise is expected in the sessions performed outside the MRI scanner, we considered the sessions OUT_1_ and OUT_2_ to act as a gold standard of measured stretch-evoked responses that is reflective of the true LLR responses, useful for comparison with the LLR-related activity measured during sessions IN_1_ and IN_2_.

EMG recorded during the two OUT sessions was processed using a standard pipeline that used all steps described above, but only used the STD filter (no expected signal from REF electrodes).

The EMG recorded during the sessions performed inside the MRI scanner was processed using three different pipelines to compare the novel processing scheme presented in this paper with two standard methods. The first method (STD) relies on the assumption that MRI-related movement artifacts are negligible and so it implements the same standard pipeline used to process the data recorded outside the scanner. In this way, the EMG signal *ŷ* is considered to be *ŷ* = *y* (defined in Sec. Simultaneous recording of fMRI and EMG data). The second method (SUB) compensates for MR-related movement artifacts assuming perfect match between the interference *r* measured by the REF electrodes and the true interference *w* (Fig. 1D). As such, it quantifies the EMG signal as *ŷ* = *y*−*r*. Finally, the third method fully implements the pipeline described in Sec. Simultaneous recording of fMRI and EMG data, quantifying the EMG signal as 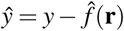.

Finally, for all sessions (*s* = [OUT_1_, IN_1_, IN_2_, OUT_2_]), filtering pipelines (*f* = [STD, SUB, ANC]), and participants (*p* = [P01, P02,…, P27]), we computed the amplitude of the stretch-evoked muscle activity *H*_*i, j,v*_|_*s, f, p*_ for all repetitions (*i* = [1, 2, …, 10]), both muscles *j* = [FCR, ECU], and all perturbations velocities (*v* = [−200, −125, −50, 50, 125, 200] deg/s) using Eq. 1. In general, averages across repetitions are indicated as 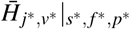 for specific levels of all other factors. The bold notation 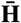 will be used to refer to the set of 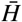 measured for all levels of one or more factors. When this operation is performed, the indices of the corresponding factors are removed (e.g. 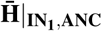 refers to the set of the mean LLR amplitudes measured during the session IN_1_ in both muscles for all participants and all perturbation velocities, when the ANC filter is used).

##### Statistical analysis

We quantified the accuracy afforded by each filtering method in identifying the true stretch-evoked muscle responses during fMRI at both the group level (combining measurements at all levels of factors muscle, velocity, participant), and at the individual perturbation level (considering each level of factors muscle, velocity, and repetition separately). With the group level analysis, we sought to quantify the agreement between the average LLR amplitude 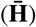 measured inside and outside the MRI for each filtering method. With the perturbation-specific analysis, we sought to quantify the deviation between each stretch-evoked response measured during MRI and the distribution of responses measured in the OUT_1_ session in the same subject, velocity, and muscle. The analysis presented below is specific to the fMRI session IN_1_; the results obtained when the following methods are applied to the session IN_2_ are reported in the Supplementary Materials.

##### Group level analysis

For the group level analysis, we used a paired Bland-Altman (BA) plot^13, 14^, a statistical method often used in chemistry^80^ and medicine^81^ to assess agreement between two measurement techniques when they are used to obtain two sets of data points in paired experimental conditions. Specifically, we compared the group level sets 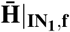 and 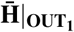 pairing the average LLR response measured for each perturbation velocity, each muscle, and for each participant. Due to the difference between responses obtained as a consequence of muscle stretch and shortening, we considering separately the subset of values measured in response to muscle stretch 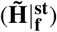 and shortening 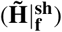.

For each of the two stimulus direction conditions (*d*) (i.e. stretch and shortening), we used BA analysis to determine bias 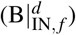 as the mean difference between paired measurements 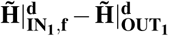, for all conditions, and the 95% limits of agreement 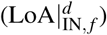, defined as the interval where given one measurement, we expect to measure the other in 95% of the cases. Each metric is measured with its own precision, expressed in terms of its 95% confidence interval, as reported in Fig. 2B. For more details on the BA analysis, please refer to^14^. Both bias and limits of agreement are expected to decrease with an increase in agreement between the two techniques. However, when the paired measurements are not obtained simultaneously with two techniques, the metrics of test-retest reliability extracted by BA analysis will be affected by both measurement error and by intrinsic physiological variability of the measurand, in this case stretch-evoked responses^12, 82^. As such, large BA or LoA measurements may be artificially inflated by physiological variability. To isolate the effects of measurement error from those of physiological variability, we thus defined our outcome measures as contrasts between the reliability measured via the IN_1_ vs. OUT_1_ comparison and the reliability measured via the OUT_2_ vs. OUT_1_ comparison.

We thus used the metrics obtained with the BA analysis to make inference on *i)* whether filtering method affects test-retest reliability, and *i)* whether filtering method affords test-retest reliability comparable with the baseline variability of the physiological process being measured (obtained for the OUT_2_ vs. OUT_1_ comparison).

We tested the null hypothesis that the estimated bias does not change when using different filtering methods by calculating the 95% confidence interval of bias for the IN_1_ vs. OUT_1_ comparison for each filtering method 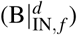, and performing three pairwise comparisons of the resulting confidence intervals (one for each pair of methods). Moreover, we tested the null hypothesis that the bias measured in the IN_1_ vs. OUT_1_ comparison was equal to the one measured in the REF comparison (OUT_2_ vs. OUT_1_) by establishing if any of the intervals 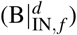 overlapped with the confidence interval 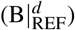 measured for the OUT_2_ vs. OUT_1_ comparison.

We conducted a similar analysis for the LoA metrics to make inference on the test-retest reliability of a specific measurement. Because the LoA is defined as an interval, test-retest reliability inferences are based on the analysis of the Jaccard index, used to quantify the relative overlap of two intervals as the ratio between their intersection and union. In our case, the Jaccard index 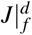 is defined as a function of the LoA measured for the comparison IN_1_ vs. OUT_1_ 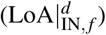, and the LoA measured for the comparison OUT_2_ vs. OUT_1_ 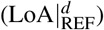, as

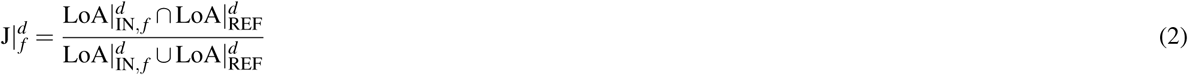

With this procedure, the test-retest error afforded by each filtering method was scaled between 0 and 1, with a score of 1 representing a perfect match of the LoA in the two conditions. Because each LoA is measured with its own precision, we determined the 95% confidence intervals for the Jaccard index using bootstrapping to numerically compute the distribution of Jaccard indices when the LoA for the comparisons IN_1_ vs. OUT_1_ and OUT_2_ vs. OUT_1_ were randomly sampled from a normal distribution with mean and standard deviation calculated from the BA analysis coefficients.

Similarly to the estimated bias analysis, we used the estimated distributions of Jaccard coefficients to make inference on whether filtering method affects test-retest reliability. Specifically, we tested the null hypothesis that there is no difference in the overlap of the limits of agreement by establishing if the 95% confidence interval of Jaccard indices overlapped for different levels of filtering method (three comparisons).

##### Perturbation-specific analysis

The goal of the perturbation-specific analysis was to quantify the departure of each stretch-evoked response measured during fMRI from the baseline values measured during the OUT_1_ session. To this aim, we computed the standardized z-score separately for each value in the set 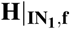 as:

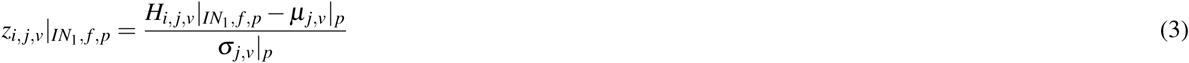

where *i, j, v, f*, and *p* are the perturbation, muscle, filtering method, velocity and participant indices, respectively; while (*µ* _*j,v*_|_*p*_) and (*σ*_*j,v*_|_*p*_) are the population mean and standard deviation of the set 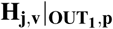. To simplify the interpretation of the results, for each stimulus direction condition *d*, we determined the standard deviation 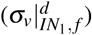 of the values in the set 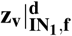. Ideally, if the elements in the sets 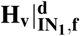 and 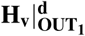 are sampled from the same normal distribution, the standard deviation 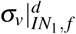 should have unitary magnitude. However, due to the physiological variability of the stretch-evoked muscle response the distribution of the elements in the two sets might have different standard deviation. As such, to have a term of comparison, we determined also the standard deviation 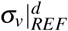 of the set of standardized scores 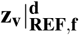 obtained applying eq. 3 to the elements in the set 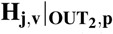.

We then tested the null hypothesis that the variance of the sets 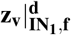 is the same for all filtering methods, performing three Bartlett tests^83^ (significance level set to p <0.05), one for each pair of sets 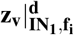,*i* = 1,2,3. Moreover, we performed three additional Bartlett tests to test the null hypothesis that the variance of the sets 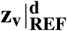 and 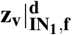 is the same for each filtering method (one test per each filtering method).

#### Statistical analysis of imaging data

We performed the analysis of all fMRI datasets using SPM12 (Wellcome Department of Cognitive Neurology, London, UK, www.fil.ion.ucl.ac.uk/spm) running on Matlab 2019a (Mathworks, Inc., Natick, MA, USA, www.mathworks.com). Functional datasets were preprocessed using a standard pipeline composed of five steps: realignment to the mean image, co-registration to structural MPRAGE, normalization to a standard MNI space template, spatial smoothing, and high-pass filtering (cut off at 128 s). For spatial smoothing, we used a Gaussian kernel with FWHM = 4 mm for voxels in the brainstem for brainstem-specific analyses (both hypothesis-testing and reliability analyses), and FWHM = 8 mm for the all other analyses. 4 mm smoothing of the brainstem was selected to account for the different expected extent of functional activations of brainstem nuclei compared to whole brain, and was used only for testing a regionally-specific (brainstem) hypotheses.

For each scanning session, we performed two first-level analyses to determine activation in the whole brain and in the brainstem, separately. While we used the same general linear model (GLM) for both analyses, we convolved it with two different hemodynamic response functions (HRF) to account for different hemodynamic response dynamics expected in different brain regions. A brainstem-specific HRF^84^ (delay of the peak = 4.5 s, delay of the undershoot = 10 s, dispersion of response = 1 s, dispersion of undershoot = 1 s, ratio of peak to undershoot = 15, length of kernel = 32 s) was used to assess activation in the brainstem, while a standard HRF (delay of the peak = 6 s, delay of the undershoot = 16 s, dispersion of response = 1 s, dispersion of undershoot = 1 s, ratio of peak to undershoot = 6, length of kernel = 32 s) was used to assess activation in all other regions. The neural response was modeled as:

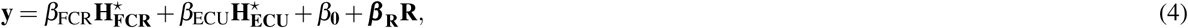

where 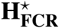 and 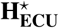 are obtained by convolution of rectangular functions (duration: 50 ms, onset: 50 ms after perturbation start, amplitude: 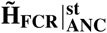 and 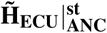, respectively) with the appropriate HRF (Fig. S6 in the Supplementary Material) to account for BOLD signal associated with LLRs of specific muscles. An additional set of 6 nuisance regressors **R** was included to account for variance in the measured signal associated with 3D head movements (translation and rotation).

To determine voxels whose BOLD signal was significantly associated with LLRs of specific muscles, we first calculated the statistical maps for the contrasts *β*_FCR_ > 0 and *β*_ECU_ > 0 for each subject, and then used the contrast images as input for the second level analysis used to determine a group level effect.

Because participants are asked to contract their muscles prior to each perturbation, there is an intrinsic statistical association between the muscle state before the perturbation and the perturbation itself. As such, given the small temporal resolution of fMRI, it is possible that some of the variance in the BOLD signal explained by LLR-related regressors might actually arise from the neural activity associated with the background contraction. To rule out this concern, we conducted two alternative analyses: one based on simulated data, and one based on the implementation of a different GLM to account for neural signal associated with background muscle activity. A detailed description of these analyses and the respective discussion and included in the Supplementary Materials.

#### Test-retest reliability of neural activations

##### Indices of reliability

Reliability of the activation maps was quantified at both the individual subject and group levels. For both levels of analysis we quantified reliability for the whole brain and for one bilateral and eight unilateral anatomical Regions of Interest (ROIs) known to be associated with the execution of motor actions: primary motor cortex (M1), primary somatosensory cortex (S1), premotor cortex (PM), superior parietal lobule (SPL), intraparietal sulcus (IPS), cerebellum (Cr), thalamus (Th), putamen (Pt), and brainstem (Bs). For each unilateral ROI, we considered two separate ROIs that refer to the left and right hemispheres. The specific brain areas included in each ROI are reported in Table 4 and were obtained by thresholding the Juelich, Harvard Subcortical, or Cerebellar MNI152 brain atlases at a probability of 50%.

**Table 4.** Definition of ROIs for test-retest reliability analysis.

| ROI |  | Brain areas |  |
| --- | --- | --- | --- |
| Name | Acronym | Name | Atlas |
| Cortical ROIs |  |  |  |
| Primary Motor Cortex | M1 | BA4a, BA4b | Juelich Atlas |
| Premotor cortex | PM | BA6 |  |
| Primary somatosensory cortex | S1 | BA1, BA2, BA3a, BA3b |  |
| Superior Parietal Lobule | SPL | BA5ci, BA5l, BA5m, BA7a, BA7m, BA7p, BA7pc |  |
| Intraparietal sulcus | IPS | hIP1, hIP2, hIP3 |  |
| Cerebellar ROI |  |  |  |
| Cerebellum | Cr | Crus 1, Crus2<br>CR I-VI, VIII, IX | Cerebellar MNI152 atlas |
| Subcortical ROIs |  |  |  |
| Thalamus | Th |  | Harvard Subcortical Atlas |
| Putamen | Pt |  |  |
| Brainstem | Bs |  |  |

For each of the brain regions listed above, with the exception of the Brainstem, we created two different ROIs that refer to the left and right hemisphere, separately. BA: Brodmann Area, CR: Cerebellar Region, hIP: human IntraParietal Area

To quantify spatial congruence between the thresholded t-maps (p <0.001, uncorrected) obtained for the two sessions IN_1_ and IN_2_, we used the Sørensen-Dice index^50, 85, 86^:

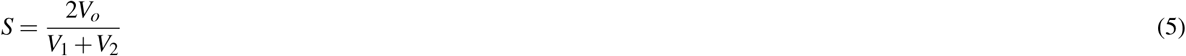

where *V*_*o*_ represents the number of supra-threshold voxels contained in both IN_1_ and IN_2_, while V_1_ and V_2_ represent the number of supra-threshold voxels contained in IN_1_ and IN_2_, respectively. This index ranges between 0 and 1, with 0 meaning no overlap and 1 meaning perfect overlap. Activation maps resulting from the first- and second-level analyses have been used to calculate the Sørensen-Dice index at the individual subject (S_*p*_) and at the group levels (S_*g*_), respectively.

While the Sørensen-Dice index is commonly used to quantify test-retest reliability of pair of fMRI-based activation maps^43, 44, 50^, the operation of thresholding used to calculate it loses a lot of information that could be relevant to quantify test-retest reliability. Specifically, a specific voxel with comparable activation in the two sessions might pass the threshold in one session but not in the other leading to an underestimate the similarity between the two maps. An opposite outcome can happen when, after thresholding, maps have similar activation regions, with the peaks of activity that, however, are located in different regions, leading to an overestimate of the reliability. To overcome these limitations, we complemented the Sørensen-Dice index with a second index, the intraclass correlation coefficient (ICC)^42, 43, 87, 88^, which instead quantifies the reliability based on the t-scores associated with each voxel, without applying any thresholding. We performed two different analyses to determine the ICC both at the individual subject (ICC_*p*_) and at the group levels (ICC_*g*_). For the individual subject analysis, we calculated the ICC applying the definition of the ICC(3,1) proposed by Shrout and Fleiss^87^ to the subset of voxels contained in a specific ROI (Tab 4) that, in the case of only two repeated measurements, simplify to:

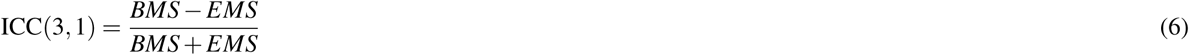

where BMS is the Between voxel Mean Square variance and EMS is the Error Mean Square variance obtained using a two-way mixed effect ANOVA, with voxel as a random effect, and session as a fixed effect.

For the group level analysis, we calculated reliability maps applying Eq. 6 to each voxel in the brain assuming subject as a random effect and session as a fixed effect^42^. To obtain the ROI-specific ICC_*g*_, we have then computed the median of the ICC distributions within each region^42^. The ICC usually ranges between 0 (low reliability) and 1 (perfect reliability). In the present study, ICCs were classified as *excellent* if they were greater than 0.75, *good* if comprised between 0.59 and 0.75, *fair* if between 0.40 and 0.58 and *poor* if smaller then 0.40, as proposed by^41^.

For each of the two subject-specific reliability indices (S_*p*_ and ICC_*p*_), we performed statistical inference to test two separate null hypotheses: *h*_0_|^1^) within each ROI, the means of the reliability indices obtained for the two regressors (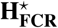 and 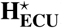) are equal; *h*_0_ |^2^) for each regressor, the means of the reliability indices obtained for the unilateral ROIs are equal in the two hemispheres. To test *h*_0_|^1^, for each of the two indices, we performed 18 paired t-tests (one per ROI plus one for the whole brain). To test *h*_0_|^2^, for each of the two indices, we performed 16 paired t-tests (two per each tested ROI).

#### Neural correlates of LLRs

##### Brainstem-specific analysis

We conducted our primary fMRI analysis to test the hypothesis that BOLD signal in any voxel in the brainstem was significantly associated with LLRs for flexors and/or extensor muscles under stretch. We used FWHM = 4 mm and the brainstem-specific HRF for this analysis. Given the regionally-specific nature of our primary hypothesis, we only included signal from brainstem voxels and applied a small-volume correction^15^,^16^ that uses random field theory to correct for the Family-Wise Error rate (FWE) for voxel-specific t-tests at the *α <* 0.05 level in the region-of-interest (number of voxels at the 2-mm isotropic resolution: 3,314). For this analysis, we combined data measured in sessions IN_1_ and IN_2_ by concatenating time-series measured in the two sessions, and adding a separate regressor to model the factor session.

##### Whole-brain analysis

We conducted a secondary analysis to test the hypothesis that BOLD signal in any voxel of the brain was significantly associated with LLRs for flexors and/or extensor muscles. For this analysis, we used FWHM = 8 mm for all voxels and the standard HRF. Given the non-specific nature of this secondary hypothesis (number of voxels tested at the group level: 228,365), we controlled for Family-wise Error rate (FWE) in the whole brain using random field theory^90, 91^ to achieve *α <* 0.05 for all voxels in the brain and applied a cluster correction of *k* = 10 (only clusters of activation larger than *k* voxels are accounted).

##### Timeseries analysis

We first extracted subject-specific *t*-scores from the first level analysis quantifying activation in the center of the two most significant clusters at the group level (MNI coordinates: [8, -42, 40] mm for the right medulla, and [-8, -30, 26] mm for the left pons), then selected individuals with the greatest difference in *t*-scores for FCR-specific activation (right medulla), and ECU-specific activation (left pons) – subj 7 and 18 respectively – and individuals with similar *t*-scores for the two regressors – subjects 9 (right medulla) and 8 (left pons) (Fig. 5). Then, we quantified the residual signal not explained by nuisance regressors (FCR residuals, nuisance regressors: ECU + head movements, ECU residuals, nuisance regressors: FCR + head movements) to evaluate muscle-specific responses of the BOLD signal. We finally segmented and averaged together the residual BOLD signal corresponding to multiple perturbations stretching specific muscles. BOLD residuals are overlaid with the regressor of interest from model 1 (scaled by the estimated *β*_*i*_ coefficient), in Fig. 5, and with scaled regressors from model 2 in Fig. S11 in the supplementary materials. Background regressor in Fig. 5 has arbitrary amplitude and is only reported for visual comparison.

## Notes

### Competing Interest Statement

AZ and FS are co-inventors of a provisional patent application on the method for EMG acquisition during fMRI.

