## Supplementary Materials for "Measurement of stretch-evoked brainstem function using fMRI"

### Behavioral Results

Of the 27 participants recruited for this study, 25 (12 for Experiment 1 and 13 for Experiment 2) completed the full experimental protocol. Two participants only completed one of the two fMRI sessions because of discomfort during imaging. For this reason, they were excluded from the all imaging analyses.

Due to technical issues affecting the quality of data collected during the experiments, other exclusions were required. Because of faulty EMG recordings during fMRI, data collected from three participants (all involved in Experiment 2) were excluded from all the analyses described above. Moreover, because of errors in slice prescription, imaging data collected in three other participants (all involved in Experiment 1) were excluded from the fMRI analyses (Sec. Test-retest reliability of neural activations and Sec. Neural correlates of LLRs).

As such, the EMG validation analysis was performed on data collected from $n = 12$ individuals, while the fMRI analyses were performed on $n = 18$ participants (9 participants exposed to Experiment 1, 9 participants exposed to Experiment 2).

Participants were able to achieve cued contraction states for both reference isometric contractions (used for normalization of filtered EMG data), and for muscle pre-conditioning (used to trigger robotic perturbations). The mean background torque measured during the set of reference isometric contractions, applied in flexion and extension prior to the beginning of each perturbation session, was (mean $\pm$ stdev) 482 $\pm$ 21mNm and 486 $\pm$ 19 mNm respectively, for session $OUT_1$, 486 $\pm$ 29 mNm and 482 $\pm$ 20 for sessions $IN_1$ and $IN_2$, 498 $\pm$ 22 mNm and 484 $\pm$ 19 mNm for session $OUT_2$. The mean filtered EMG amplitude measured during the same isometric contractions in the FCR and ECU was 24.61 $\pm$ 15.18 $\mu$V and 91.89 $\pm$ 40.26 $\mu$V respectively, for session $OUT_1$, 36.23 $\pm$ 34.99 $\mu$V and 99.57 $\pm$ 48.48 $\mu$V for sessions $IN_1$ and $IN_2$, and 27.19$\pm$19.64 $\mu$V and 90.70 $\pm$ 29.04$\mu$V for session $OUT_2$.

The mean background torque measured prior to each perturbation (average between measurements during flexion and extension) was 183 $\pm$ 19 mNm, 192 $\pm$ 20 mNm, 191 $\pm$ 18 mNm, 187 $\pm$ 18 mNm, respectively for $OUT_1$, $IN_1$, $IN_2$, and $OUT_2$, with the time required to trigger a robotic perturbation equal to 1.55 $\pm$ 0.68 s, 2.90 $\pm$ 1.75 s, 2.78 $\pm$ 1.55 s, 1.44 $\pm$ 0.62 s, respectively for the four sessions. The mean time elapsed between two consecutive perturbations was 10.12 $\pm$ 1.43 s, 14.08 $\pm$ 3.03 s, 14.11 $\pm$ 2.40 s, 9.88 $\pm$ 1.36 s. The group level distributions of the parameters listed above are shown in the Fig. S1A-B.

The mean timeseries of the processed EMG recorded during each session of the experiment 1 and processed using the three different filtering pipelines are reported in Figs. S7 and S8 in the supplementary materials, respectively for data measured in Experiment 1 and Experiment 2.

### Supplementary Analyses

The GLM selected to partition the variance of the measured BOLD signal (Eq. 4 of the main manuscript) does not include separate regressors for background activity and LLR amplitude. Because participants are asked to contract their muscles prior to each perturbation, there is an intrinsic statistical association between the muscle state before the perturbation and the perturbation itself. As such, given the poor temporal resolution of fMRI, it is possible that some of the variance in the BOLD

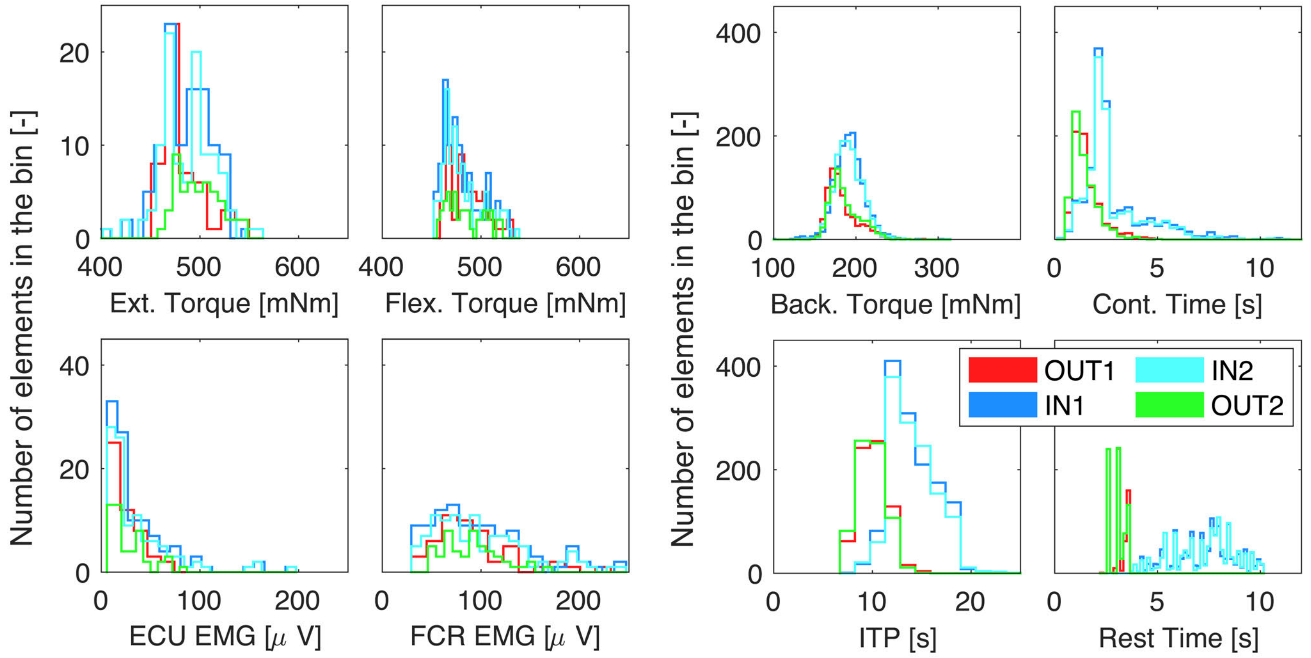


**Figure S1.** (**A**) Distribution of the behavioral parameters measured during the reference isometric contraction applied prior to the beginning of each session and utilized for the normalization of the stretch-evoked EMG recordings. (**B**) Distribution of the perturbation-specific behavioral parameters.

signal explained by LLR-related regressors might actually arise from the neural activity associated to background contraction. To rule out this concern, we conducted two analyses, one based on simulated data, and one based on the implementation of a different GLM on the fMRI data collected in Experiment 2.

### Model-based analysis

We initially sought to quantify whether the implemented analysis method is sufficiently sensitive and specific in identifying neural activity related to LLR in presence of correlated background activity. To this goal we have conducted a simulated signal analysis following the methods presented in our previous work[1].

**Methods** We simulated nine different representative brain voxels $V_1$ to $V_9$ that selectively respond only to linear combinations of conditions of flexor and extensor muscle activity, defined in terms of LLR events ($\mathbf{H}^{\star}_{\mathbf{F}}$ and $\mathbf{H}^{\star}_{\mathbf{E}}$ , respectively for flexor and extensor) and background contraction ($\mathbf{B}^{\star}_{\mathbf{F}}$ and $\mathbf{B}^{\star}_{\mathbf{E}}$, respectively for flexor and extensor). Specifically, $\mathbf{H}^{\star}_{\mathbf{F}}$ and $\mathbf{H}^{\star}_{\mathbf{E}}$ are obtained by convolution of rectangular functions (duration: 50 ms, onset: 50 ms after perturbation start, amplitude: $\tilde{\mathbf{H}}_{\mathbf{FCR}}|^{\mathbf{st}}_{\mathbf{ANC,P4}}$ and $\tilde{\mathbf{H}}_{\mathbf{ECU}}|^{\mathbf{st}}_{\mathbf{ANC,P4}}$, respectively) with the standard HRF (Fig. S6 in the Supplementary Material). Similarly, the terms $\mathbf{B}^{\star}_{\mathbf{F}}$ and $\mathbf{B}^{\star}_{\mathbf{E}}$ are given by the convolution of rectangular functions (unitary magnitude in the time interval when the measured torque has a magnitude greater than 25% of the desired torque) with the standard HRF (Fig. S6 in the Supplementary Material). To simulate all four conditions, we used the EMG and torque data measured in a representative participant (participant P4). A schematic representation of the the simulated voxels is reported in Tabs. S1 and S2.

**Table S1.** Definition of voxel used in the simulation analysis

| Effects | Flexor | Extensor | Both |
|---|---|---|---|
| $\mathbf{H}^{\star}$ | $V_1$ | $V_2$ | $V_3$ |
| $\mathbf{B}^{\star}$ | $V_4$ | $V_5$ | $V_6$ |
| $\mathbf{H}^{\star}+\mathbf{B}^{\star}$ | $V_7$ | $V_8$ | $V_9$ |

For each of the simulated voxels, we then added zero-mean Gaussian noise and down-sampled the signal with a period $T_R$ = 1.225 s, to simulate realistic BOLD measurements specific to either cortical voxels (Signal-To-Noise ratio equal to 1.2) or brainstem voxels (Signal-to-Noise ratio equal to 0.5). Finally, we quantified the sensitivity that the two proposed GLMs (GLM$_1$

in Eq. 4 and $GLM_2$ in Eq. S1) have in identifying the four different regressors, by computing the true positive (TPR) and false positive (FPR) rates afforded when either of the two GLM is used to fit the signal simulated in each of the nine different voxels.

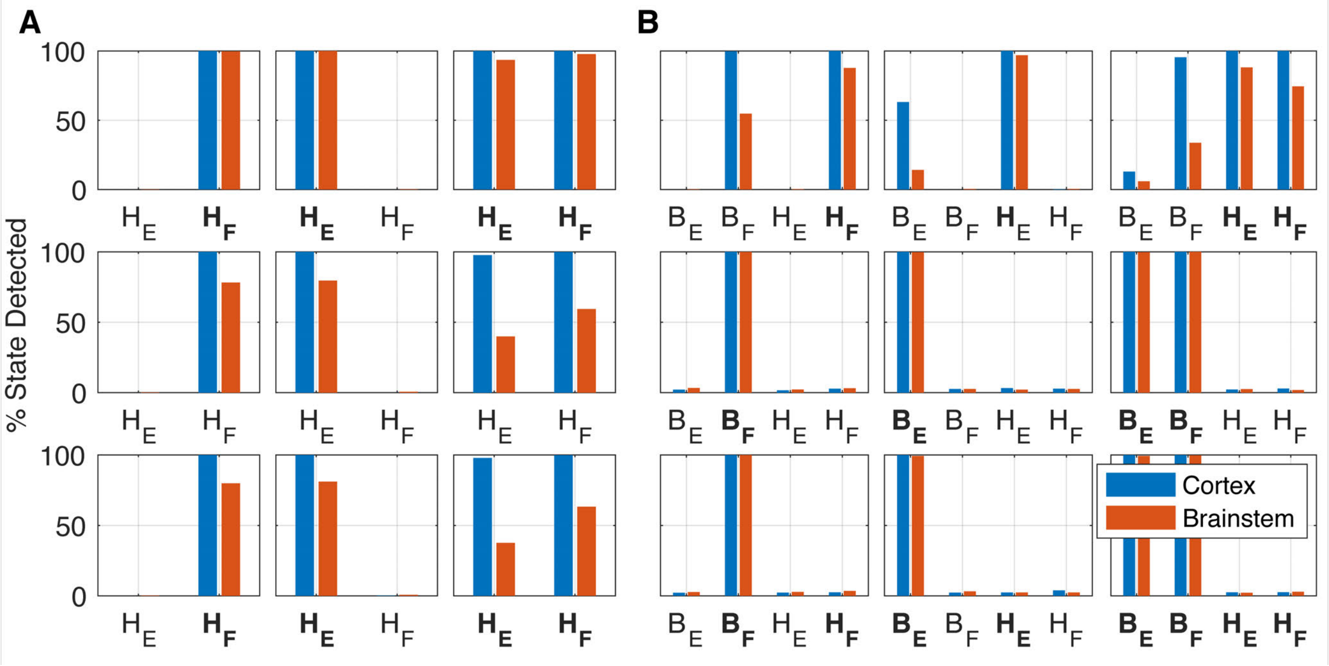


**Figure S2.** Identification rate for $GLM_1$ (**A**) and $GLM_2$ (**B**). Bold labels indicate true positive association between the indicated regressor and the BOLD signal measured in the corresponding voxel.

**Results** The results show that the model $GLM_1$ affords the highest TPR for LLR-specific voxels in both cortical and brainstem voxels, and for both flexor and extensor regressors ($V_1$ and $V_2$, respectively; Fig. S2A, top row and Tab. S2). However, it also returns high FPR for LLR-related activity in voxels whose activity is associated to the application fo background torque (Fig. S2A, middle and Tab. S2), though the FPR is smaller in the brainstem voxels than in the cortical ones.

The inclusion of the background-specific regressors drastically reduces the FPR of LLR-related activity in background-specific voxels for both flexor and extensor ($V_4$ and $V_5$, respectively; Fig. S2B, middle row and Tab. S2). However, while TPR remains high in LLR-specific voxels (Fig. S2A, top row and Tab. S2), it decreases to almost zero in voxels that responds to both LLR and background activity ($V_7$ and $v_8$; Fig. S2A, bottom row and Tab. S2).

### Alternative GLM analysis

We conducted an alternative statistical analysis for the BOLD fMRI data that aims to determine the LLR-related neural activity including a correction for the possible confounding effect introduced by the background-related activity.

**Methods** Methods closely resembled those described in Sec. Statistical analysis of imaging data for most aspects. Separately for each session, we performed two first-level analyses to determine activation in the whole brain and in the brainstem. While we used the same general linear model (GLM) for the whole-brain and brainstem analyses, we used regionally-specific hemodynamic response functions (HRF) for different brain regions. A brainstem-specific HRF[2] was used to assess activation in the brainstem, while a standard HRF was used to assess activation in all other regions. In this alternative GLM ($GLM_2$) analysis, the neural response was modeled as:

$$\mathbf{y} = \beta_{\mathrm{FCR,L}}\mathbf{H}^{\star}_{\mathbf{FCR}} + \beta_{\mathrm{ECU,L}}\mathbf{H}^{\star}_{\mathbf{ECU}} + \beta_{\mathrm{FCR,B}}\mathbf{B}^{\star}_{\mathbf{FCR}} + \beta_{\mathrm{ECU,B}}\mathbf{B}^{\star}_{\mathbf{ECU}} + \beta_{\mathbf{0}} + \boldsymbol{\beta}_{\boldsymbol{R}}\mathbf{R} \tag{S1}$$

where $\mathbf{H}^{\star}_{\mathbf{FCR}}$ and $\mathbf{H}^{\star}_{\mathbf{ECU}}$ are obtained by convolution of rectangular functions (duration: 50 ms, onset: 50 ms after perturbation start, amplitude: $\tilde{\mathbf{H}}_{\mathbf{FCR}}|^{\mathbf{st}}_{\mathbf{ANC}}$ and $\tilde{\mathbf{H}}_{\mathbf{ECU}}|^{\mathbf{st}}_{\mathbf{ANC}}$, respectively) with the appropriate HRF (Fig. S6 in the Supplementary Material). The terms $\mathbf{B}^{\star}_{\mathbf{FCR}}$ and $\mathbf{B}^{\star}_{\mathbf{ECU}}$ are now added to the model to account for metabolic activity associated with muscle-specific contraction state. They are similarly obtained convolving rectangular functions (unitary magnitude in the time interval when the measured torque is in magnitude greater than 25% of the desired torque) with the appropriate HRF (Fig. S6 in the Supplementary Material). As in all analyses, a set of 6 nuisance regressors **R** was included to account for variance in the measured signal associated with the 3D head movements (translation and rotations).

**Table S2.** Percent state detected in the different voxels for the two GLMs. Bolded labels represents true positive identification rates, while un-bolded labels represents false positive identification rate

| Voxel | Effects | $H_F$ | $H_E$ | $B_F$ | $B_E$ | $H_F$ | $H_E$ | $B_F$ | $B_E$ |
|---|---|---|---|---|---|---|---|---|---|
| **GLM$_1$** | | Cortex | | | | Brainstem | | | |
| $V_1$ | $\mathbf{H_F^\star}$ | **1** | 0 | - | - | **0.99** | 0.01 | - | - |
| $V_2$ | $\mathbf{H_E^\star}$ | 0 | **1** | - | - | 0.01 | **1** | - | - |
| $V_3$ | $\mathbf{H_F^\star} + \mathbf{H_{ECU}^\star}$ | **1** | **1** | - | - | **0.97** | **0.93** | - | - |
| $V_4$ | $\mathbf{H_E^\star}$ | 1 | 0 | - | - | 0.78 | 0.01 | - | - |
| $V_5$ | $\mathbf{B_E^\star}$ | 0 | 1 | - | - | 0.01 | 0.79 | - | - |
| $V_6$ | $\mathbf{B_F^\star} + \mathbf{B_{ECU}^\star}$ | 0.99 | 0.97 | - | - | 0.59 | 0.39 | - | - |
| $V_7$ | $\mathbf{H_F^\star} + \mathbf{B_F^\star}$ | **1** | 0 | - | - | **0.79** | 0.01 | - | - |
| $V_8$ | $\mathbf{H_E^\star} + \mathbf{B_E^\star}$ | 0.001 | **1** | - | - | 0.01 | **0.81** | - | - |
| $V_9$ | $\mathbf{H_F^\star} + \mathbf{H_E^\star} + \mathbf{B_F^\star} + \mathbf{B_E^\star}$ | **0.99** | **0.97** | - | - | **0.63** | **0.37** | - | - |
| **GLM$_2$** | | Cortex | | | | Brainstem | | | |
| $V_1$ | $\mathbf{H_F^\star}$ | **1** | 0 | 1 | 0 | **0.88** | 0 | 0.55 | 0 |
| $V_2$ | $\mathbf{H_E^\star}$ | 0 | **1** | 0 | 0.63 | 0 | **0.97** | 0 | 0.14 |
| $V_3$ | $\mathbf{H_F^\star} + \mathbf{H_E^\star}$ | **1** | **1** | 0.95 | 0.13 | **0.74** | **0.88** | 0.34 | 0.06 |
| $V_4$ | $\mathbf{B_F^\star}$ | 0.03 | 0.02 | **1** | 0.02 | 0.03 | 0.02 | **1** | 0.03 |
| $V_5$ | $\mathbf{B_E^\star}$ | 0.03 | 0.03 | 0.03 | **1** | 0.03 | 0.02 | 0.03 | **1** |
| $V_6$ | $\mathbf{B_F^\star} + \mathbf{B_E^\star}$ | 0.03 | 0.02 | **1** | **1** | 0.02 | 0.03 | **1** | **1** |
| $V_7$ | $\mathbf{H_F^\star} + \mathbf{B_F^\star}$ | **0.03** | 0.02 | **1** | 0.02 | **0.04** | 0.03 | **1** | 0.03 |
| $V_8$ | $\mathbf{H_E^\star} + \mathbf{B_E^\star}$ | 0.04 | **0.02** | 0.02 | **1** | 0.02 | **0.02** | 0.03 | **0.99** |
| $V_9$ | $\mathbf{H_F^\star} + \mathbf{H_E^\star} + \mathbf{B_F^\star} + \mathbf{B_E^\star}$ | **0.03** | **0.03** | **1** | **1** | **0.03** | **0.02** | **1** | **0.99** |

Bolded number represents true positive identification rates, while un-bolded numbers represents false positive identification rates

**Results** For the brainstem-specific analysis, muscle-specific maps included several clusters of activation associated only to the LLR-specific regressors (Fig. S3, Tab. S3). In fact, no supra-threshold voxels were identified to correlate with the background-specific regressors (Fig. S4). LLR-related activity specific to FCR included a total of 18 voxels, including two clusters spanning bilaterally the midbrain, two clusters spanning bilaterally the pons, and two clusters in the right superior medulla (Fig. S3 top, and Table S3). LLR-related activity specific to ECU included a total of 16 voxels, including one voxel in the right midbrain, four clusters spanning bilaterally the pons, and one cluster in the right medulla (Fig. S3 top, and Table S3).

In contrast to what we observed in the brainstem-specific analysis, in the whole brain analysis muscle-specific maps included several clusters of activation associated to both LLR-specific (Fig. S3 bottom, Tab. S4, and Tab. S5) and background-specific regressors (Fig. S3 bottom, Tab. S6, and Tab. S7). Specifically, cortical activation associated to the two background-specific regressors ($\mathbf{B_{FCR}^\star}$ and $\mathbf{B_{ECU}^\star}$) primarily included overlapped clusters of activation in the contralateral sensorimotor cortex and inferior parietal lobule, in the ipsilateral insular and opercular cortex, and in the bilateral visual cortex. Cerebellar activity included overlapped clusters in the ipsilateral region V and contralateral region VI, while in the subcortical regions activity was primarily detected in the contralateral putamen (Fig. S3 bottom, Tab. S6, and Tab. S7). LLR-related cortical activity specific to FCR primarily included contralateral activation in the sensorimotor cortex, inferior parietal lobule and bilateral activation in the visual cortex. Cerebellar activity could be instead observed bilaterally in the regions I-IV and VIII, and contralaterally in the region VI and in the Crus II, while subcortical activity could be observed in the contralateral putamen and thalamus and in the ipsilateral caudate nucleus (Fig. S3 bottom, Tab. S4). As for the LLR-related activity specific to ECU, no subcortical activity could be observed, with cortical activity observable only in the ipsilateral visual cortex. Cerebellar activity was instead more diffuse and included bilateral activation in the region VIII and Crus I, ipsilateral activity in the regions V and X, and contralateral activity in the region VI and in the Crus II (Fig. S3 bottom, Tab. S4).

### Discussion

The results of the simulation analysis show that voxels that are positively associated with the regressor $\mathbf{H_F^\star}$ in GLM$_1$ might respond to LLR activity, background activity, or both–i.e. $V_1$, $V_4$, or $V_7$, respectively (Fig. S2A, first column). On the contrary, when using GLM$_2$, voxels that are positively associated with the regressor $\mathbf{H_F^\star}$ can only respond to LLR-related activity–i.e. $V_1$–since a 0% positive identification rate characterize voxels $V_4$ and $V_7$. For all three voxels, however, the background-specific regressor $\mathbf{B_F^\star}$ dominates, reducing the specificity of the regressor $\mathbf{B_F^\star}$ in voxels that only respond to LLR activity ($V_1$) and the

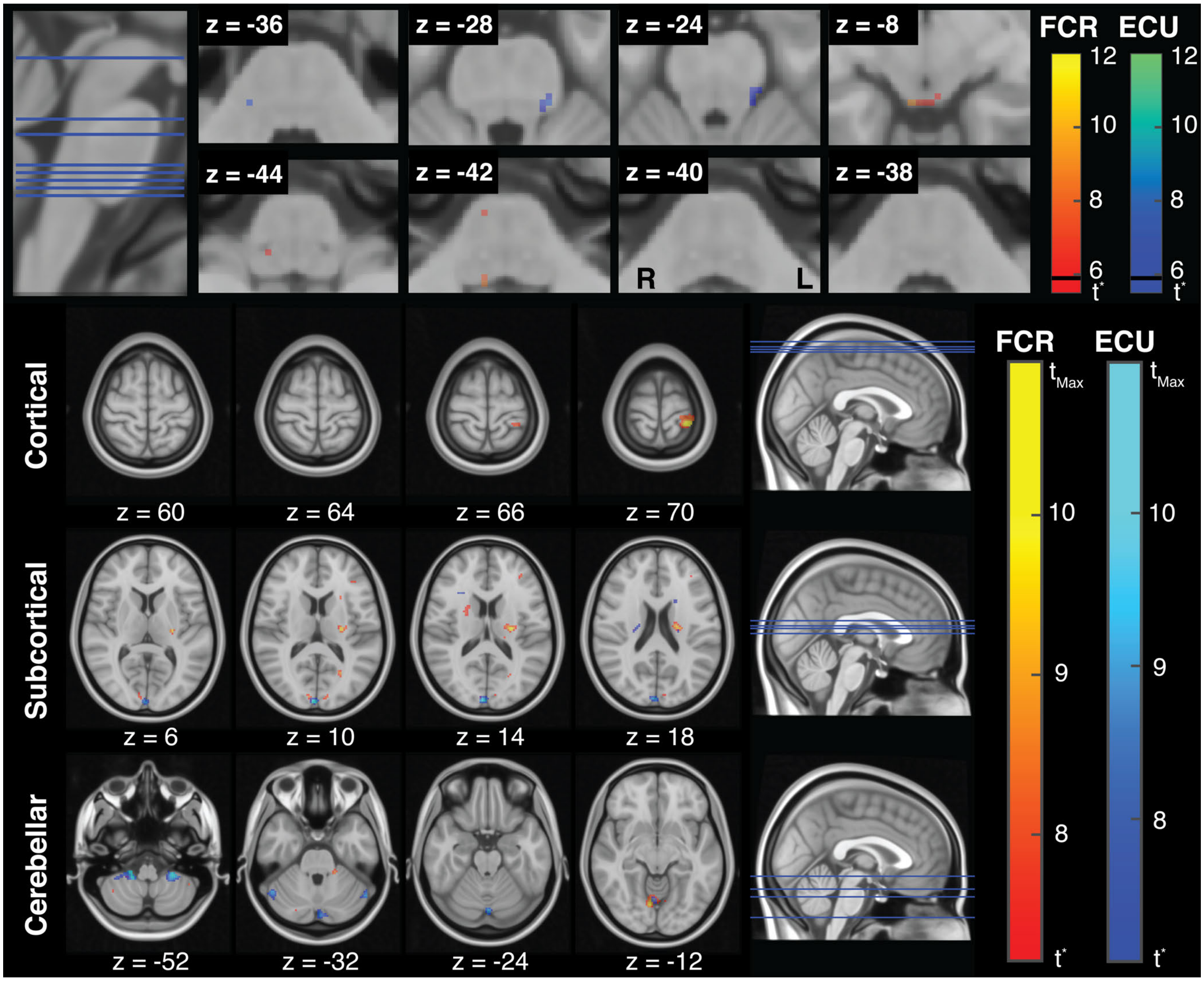


**Figure S3.** (**Top**) LLR-specific activation maps in the brainstem for FCR and ECU. The maps refer to contrasts $\beta_{\text{FCR,L}} > 0$ and $\beta_{\text{ECU,L}} > 0$ obtained for the brainstem-specific analysis. The threshold t-statistic, obtained after FWE correction, was equal to 5.62 for both regressors. For reference, the t-statistic that refer to a Bonferroni correction ($t_{\text{Bon}}$= 5.8) is marked on each colorbar with a black line. (**Bottom**) LLR-specific activation maps in the whole-brain for FCR and ECU. The threshold t-statistic, obtained after FWE correction, was set to 7.28 and 6.99, respectively for FCR and ECU, while $t_{Max}$ was equal to 12.81 and 11.95, respectively. Colorbars are saturated at t = 10 for better visualization of t-statistic gradients. All statistical parametric maps are overlaid on axial slices of the standard Montreal Neurological Institute 152 template, with reported z coordinate in mm

sensitivity of the regressor $\mathbf{H}^{\star}_{\mathbf{F}}$ in voxels that instead respond to both LLR and background activity ($V_7$), for both cortical and brainstem voxels.

The same observation can be made considering voxels that respond to activity in the extensors. Therefore, it is possible to conclude that when using the model $GLM_2$, background related activity will always be identified if present, and dominate over the LLR-related regressor. However, the brainstem activation maps obtained using the $GLM_2$ model show the complete absence of voxels whose signal is significantly associated with the background specific regressors (Fig. S4). This evidence, together with the observation that the clusters of activations included in the LLR-specific regressors (Fig. S3, Tab. S3) are closely aligned with the clusters of activations identified by $GLM_1$ (Fig. 3, Tab. 1), suggest that the identified brainstem activity is truly associated with LLR-related activity and not with background activation. The insight from the two analyses presented above advances that it is unlikely that some of the variance in the BOLD signal explained by LLR-related regressors might arise from the neural activity associated to background contraction.

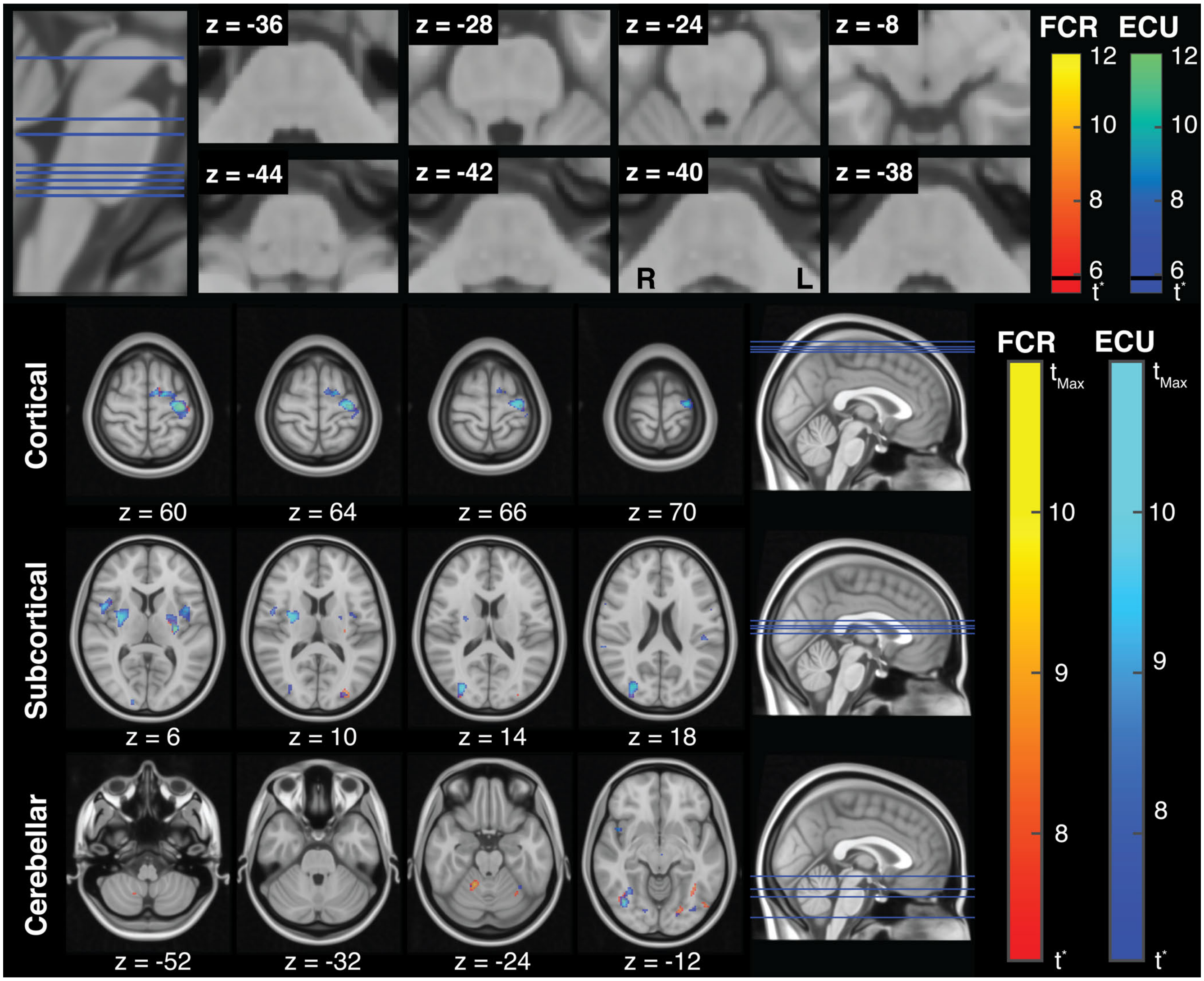


**Figure S4.** (**Top**) Activation maps in the brainstem for FCR and ECU background activity. The maps refer to contrasts $\beta_{\mathrm{FCR,B}} > 0$ and $\beta_{\mathrm{ECU,B}} > 0$ obtained for the brainstem-specific analysis. The threshold t-statistic, obtained after FWE correction, was equal to 5.62 for both regressors. For reference, the t-statistic that refer to a Bonferroni correction ($t_{\mathrm{Bon}}$= 5.8) is marked on each colorbar with a black line. (**Bottom**) Activation maps in the whole brain for FCR and ECU background activity. The threshold t-statistic, obtained after FWE correction, was set to 7.27 and 7.17, respectively for FCR and ECU, while $t_{Max}$ was equal to 11.44 and 16.66, respectively. Colorbars are saturated at t = 10 for better visualization of t-statistic gradients. All statistical parametric maps are overlaid on axial slices of the standard Montreal Neurological Institute 152 template, with reported z coordinate in mm

In contrast to the similarity observed between the results obtained for the two GLMs in the brainstem-specific analysis, at the whole brain level the two GLMs produce fairly different results. Since both LLRs and isometric contractions are intrinsic motor actions, they are expected to share most of the same cortical substrates. As such, as predicted by the model, it is possible to observe a decrease of the sensitivity that the model has in identifying LLR-related activity, mostly evident in ECU-specific maps (Fig. S3 bottom, Tab. S4).

In summary, due to the co-linearity between the LLR-specific and background-specific regressors–i.e. after the application of a background torque there is always a perturbation that elicits a reflex response in the pre-conditioned muscles–our experimental protocol does not afford enough statistical power to reliably decouple the LLR-related and background-related activity at the cortical level. However, as demonstrated by the interaction between the results of of the simulation analysis and the alternative GLM analysis, it is very likely that the measured activations are specific to LLR-related responses, and minimally affected by statistically associated activity arising from background contraction.

**Table S3.** Significance table for the brainstem-specific analysis

| **Brainstem Region** | | **Muscle** | **Peak-Level** | | **Cluster-Level** | | **Peak coordinate** [mm] | | |
|---|---|---|---|---|---|---|---|---|---|
| Area | Laterality | | $t_{max}$ | $p_{FWE}$ | Voxels in cluster | $p_{FWE}$ | x | y | z |
| Midbrain | R | FCR | 8.86 | <0.001 | 11 | <0.001 | 6 | -36 | -6 |
| | L | FCR | 5.84 | 0.02 | 1 | 0.032 | -4 | -38 | -16 |
| | R | ECU | 6.13 | 0.018 | 1 | 0.018 | 8 | -32 | –16 |
| Pons | L | FCR | 6.77 | 0.005 | 2 | 0.007 | -10 | -26 | -32 |
| | R | FCR | 5.97 | 0.024 | 1 | 0.018 | 8 | -22 | -40 |
| | R | ECU | 7.20 | 0.002 | 8 | <0.001 | 16 | -30 | -32 |
| | L | ECU | 7.20 | 0.002 | $11^{a}$ | <0.001 | -12 | -32 | -26 |
| | L | ECU | 6.57 | 0.008 | 2 | 0.008 | -8 | -34 | -44 |
| | L | ECU | 6.21 | 0.015 | - part of Cluster *a* - | | -10 | -32 | -22 |
| Medulla | R | FCR | 8.09 | <0.001 | 2 | 0.007 | 8 | -44 | -40 |
| | R | FCR | 5.64 | 0.047 | 1 | 0.018 | 10 | -36 | -42 |
| | R | ECU | 6.10 | 0.019 | 2 | 0.008 | 8 | -40 | -50 |

Superscripts on the cluster volume, when present, refer to clusters with multiple peaks. Coordinates are express in MNI space.

**Table S4.** Significance table for the whole-brain analysis for FCR-specific LLR-related activation

| **Region** | | **Peak-Level** | | **Cluster-Level** | | **Peak coordinate** [mm] | | |
|---|---|---|---|---|---|---|---|---|
| Area | Laterality | $t_{max}$ | $p_{FWE}$ | Voxels in cluster | $p_{FWE}$ | x | y | z |
| **Cortical Regions** | | | | | | | | |
| Sensorimotor Cortex | L | 12.82 | <0.001 | $135^{a}$ | <0.001 | -32 | -38 | 72 |
| | L | 9.98 | 0.002 | - part of Cluster *a* - | | -24 | -30 | 76 |
| Lingual Gyrus | R | 10.77 | 0.001 | $377^{b}$ | <0.001 | 8 | -70 | -8 |
| Visual Cortex | R | 9.18 | 0.004 | - part of Cluster *b* - | | 8 | -92 | 12 |
| | R | 9.12 | 0.005 | - part of Cluster *b* - | | 10 | -84 | 4 |
| | L | 8.14 | 0.016 | $35^{c}$ | <0.001 | -4 | -90 | 24 |
| | R | 8.06 | 0.018 | - part of Cluster *d* - | | 8 | -88 | 28 |
| | L | 7.90 | 0.022 | - part of Cluster *c* - | | -10 | -90 | 12 |
| Inferior parietal lobule | L | 9.55 | 0.003 | 68 | <0.001 | -44 | -52 | 38 |
| Frontal Pole | R | 8.64 | 0.008 | 10 | 0.003 | 40 | 38 | 4 |
| | L | 8.21 | 0.014 | 10 | 0.003 | -36 | 40 | 18 |
| Paracingulate Gyrus | L | 8.39 | 0.011 | 20 | 0.001 | -8 | 34 | 32 |
| Cuneal Cortex | R | 8.39 | 0.011 | $75^{d}$ | <0.001 | 10 | -82 | 38 |
| **Cerebellar Regions** | | | | | | | | |
| CB I-IV | R | 9.97 | 0.002 | 105 | <0.001 | 26 | -32 | -36 |
| | L | 8.51 | 0.010 | - part of Cluster *e* - | | -18 | -36 | -30 |
| CB VI | L | 9.10 | 0.005 | $117^{e}$ | <0.001 | -24 | -38 | -36 |
| CB VIII | L | 9.05 | 0.005 | - part of Cluster *e* - | | -30 | -44 | -42 |
| | R | 8.37 | 0.012 | 65 | <0.001 | 36 | -56 | -40 |
| | R | 8.23 | 0.014 | 14 | 0.002 | 42 | -64 | -36 |
| Crus II | L | 8.00 | 0.019 | 23 | <0.001 | -32 | -60 | -40 |
| **Subcortical Regions** | | | | | | | | |
| Putamen/Thalamus | L | 10.91 | <0.001 | $115^{f}$ | <0.001 | -24 | -16 | 16 |
| | L | 9.57 | 0.003 | - part of Cluster *f* - | | -26 | -18 | 8 |
| Caudate | R | 8.04 | 0.018 | 26 | <0.001 | 24 | 0 | 18 |

Superscripts on the cluster volume, when present, refer to clusters with multiple peaks. Coordinates are expressed in MNI space.

### Limitations

The main limitation of our experimental protocol relates to the fact that the fMRI scanning sequence currently uses an Echo Planar Imaging (EPI) routinely used for fMRI studies. While the results obtained suggest that such imaging sequence has

**Table S5.** Significance table for the whole-brain analysis for ECU-specific LLR-related activation

| **Region** | | **Peak-Level** | | **Cluster-Level** | | **Peak coordinate** [mm] | | |
|---|---|---|---|---|---|---|---|---|
| Area | Laterality | $t_{max}$ | $p_{FWE}$ | Voxels in cluster | $p_{FWE}$ | x | y | z |
| **Cortical Regions** | | | | | | | | |
| Visual Cortex | R | 10.61 | 0.001 | 139[a] | <0.001 | 6 | -94 | 14 |
| | R | 8.01 | 0.012 | - part of Cluster *a* - | | 0 | -88 | 30 |
| **Cerebellar Regions** | | | | | | | | |
| CB X | R | 11.96 | <0.001 | 81[b] | <0.001 | 20 | -36 | -50 |
| CBVIII | R | 11.74 | <0.001 | - part of Cluster *b* - | | 16 | -44 | -52 |
| | L | 10.20 | 0.001 | 142[c] | <0.001 | -26 | -40 | -50 |
| | R | 7.71 | 0.019 | - part of Cluster *b* - | | 34 | -42 | -50 |
| | L | 7.39 | 0.029 | - part of Cluster *c* - | | -18 | -48 | -46 |
| | L | 7.21 | 0.037 | - part of Cluster *c* - | | -14 | -44 | -54 |
| Crus I | L | 9.58 | 0.002 | 71 | <0.001 | -52 | -52 | -34 |
| | R | 9.38 | 0.002 | 92[d] | <0.001 | 50 | -58 | -32 |
| | L | 8.86 | 0.004 | - part of Cluster *d* - | | -52 | -62 | -30 |
| | R | 8.14 | 0.010 | 15 | 0.004 | 38 | -44 | -40 |
| Crus II | L | 9.06 | 0.003 | 225[e] | <0.001 | -2 | -84 | -32 |
| Vermis IV | L | 9.04 | 0.003 | - part of Cluster *e* - | | -2 | -80 | -24 |
| CB V | R | 8.87 | 0.004 | - part of Cluster *e* - | | 4 | -66 | -8 |
| CB VI | L | 8.16 | 0.010 | 19 | 0.003 | -40 | -40 | -36 |

Superscripts on the cluster volume, when present, refer to clusters with multiple peaks. Coordinates are expressed in MNI space.

sufficient contrast-to-noise ratio and robustness from motion artifacts to detect meaningful change in both cortical ROIs and deep brainstem nuclei, we are working to include a high-resolution multi-shot sequence that has specifically been developed for brainstem imaging in order to improve the contrast-to-noise ratio to about 10-20%[3].

**Table S6.** Significance table for the whole-brain analysis for FCR-specific background-related activation

| **Region** | | **Peak-Level** | | **Cluster-Level** | | **Peak coordinate** [mm] | | |
|---|---|---|---|---|---|---|---|---|
| Area | Laterality | $t_{max}$ | $p_{FWE}$ | Voxels in cluster | $p_{FWE}$ | x | y | z |
| **Cortical Regions** | | | | | | | | |
| Sensorimotor Cortex | L | 10.59 | 0.001 | 149[a] | <0.001 | -52 | -22 | 42 |
| | L | 10.47 | 0.001 | 486[b] | <0.001 | -36 | -20 | 62 |
| | L | 9.80 | 0.002 | 44[c] | <0.001 | -54 | 4 | 36 |
| | L | 9.31 | 0.004 | - part of Cluster *b* - | | -36 | -22 | 52 |
| | L | 8.79 | 0.007 | - part of Cluster *b* - | | -36 | -10 | 54 |
| | L | 8.70 | 0.008 | 22 | 0.001 | -6 | -12 | 58 |
| | L | 8.23 | 0.014 | 25 | <0.001 | -12 | -2 | 58 |
| | L | 8.10 | 0.016 | 44[d] | <0.001 | -6 | 4 | 46 |
| | L | 8.00 | 0.019 | 14 | 0.002 | -32 | -38 | 50 |
| Visual Cortex | L | 10.19 | 0.001 | 87 | <0.001 | -50 | -72 | -6 |
| | R | 9.71 | 0.002 | 98[e] | <0.001 | 40 | -66 | -10 |
| | L | 9.43 | 0.003 | 45 | <0.001 | -28 | -86 | 12 |
| | L | 8.89 | 0.006 | 63 | <0.001 | -22 | -78 | -12 |
| | R | 8.41 | 0.011 | - part of Cluster *e* - | | 46 | -58 | 0 |
| | R | 7.81 | 0.024 | 17 | 0.001 | 32 | -88 | 20 |
| Insular Cortex | R | 9.32 | 0.004 | 93[f] | <0.001 | 36 | 10 | 4 |
| Broca's Area | L | 8.76 | 0.007 | - part of Cluster *c* - | | -56 | 8 | 28 |
| | R | 7.45 | 0.039 | - part of Cluster *e* - | | 44 | 14 | 10 |
| Inferior parietal lobule | L | 8.70 | 0.008 | - part of Cluster *a* - | | -56 | -20 | 26 |
| Operculum | R | 8.21 | 0.014 | - part of Cluster *f* - | | 44 | 4 | 0 |
| Cingulate Gyrus | L | 8.06 | 0.017 | - part of Cluster *d* - | | -6 | 2 | 38 |
| **Cerebellar Regions** | | | | | | | | |
| CB V | R | 9.36 | 0.003 | 69 | <0.001 | 16 | -50 | -24 |
| CB VIIIa | R | 8.31 | 0.012 | 15 | 0.001 | 20 | -64 | -48 |
| CB VI | L | 8.14 | 0.016 | 17 | 0.001 | -28 | -60 | -20 |
| **Subcortical Regions** | | | | | | | | |
| Putamen | L | 11.49 | <0.001 | 255[g] | <0.001 | -28 | -18 | 8 |
| Pallidum | L | 8.87 | 0.006 | - part of Cluster *g* - | | -22 | -6 | -6 |

Superscripts on the cluster volume, when present, refer to clusters with multiple peaks. Coordinates are expressed in MNI space.

**Table S7.** Significance table for the whole-brain analysis for ECU-specific background-related activation

| Region | | Peak-Level | | Cluster-Level | | Peak coordinate [mm] | | |
|---|---|---|---|---|---|---|---|---|
| Area | Laterality | $t_{max}$ | $p_{FWE}$ | Voxels in cluster | $p_{FWE}$ | x | y | z |
| **Cortical Regions** | | | | | | | | |
| Sensorimotor Cortex | L | 16.67 | 0.000 | 2066[a] | 0.000 | -8 | 4 | 36 |
| | L | 15.13 | 0.000 | - part of Cluster *a* - | | -32 | -18 | 64 |
| | L | 12.53 | 0.000 | - part of Cluster *a* - | | -34 | -20 | 46 |
| | L | 8.38 | 0.010 | 15 | 0.002 | -8 | -24 | 46 |
| | R | 7.84 | 0.020 | 18 | 0.002 | 34 | -6 | 52 |
| Insular Cortex | L | 12.07 | 0.000 | - part of Cluster *d* - | | -36 | 4 | 0 |
| | R | 11.96 | 0.000 | - part of Cluster *e* - | | 44 | 2 | 0 |
| Visual Cortex | R | 11.56 | 0.000 | 165 | 0.000 | 30 | -80 | 18 |
| | R | 10.85 | 0.001 | 255 | 0.000 | 38 | -66 | -14 |
| | R | 9.89 | 0.002 | 23 | 0.001 | 16 | -98 | 6 |
| | L | 8.08 | 0.015 | 66[b] | 0.000 | -30 | -80 | -14 |
| | L | 7.87 | 0.019 | 16 | 0.002 | -26 | -88 | 12 |
| | L | 7.84 | 0.020 | - part of Cluster *b* - | | -38 | -78 | -12 |
| Inferior parietal lobule | L | 9.15 | 0.004 | 48[c] | 0.000 | -58 | -20 | 26 |
| | L | 8.59 | 0.008 | 31 | 0.000 | -50 | -30 | 24 |
| | R | 8.58 | 0.008 | 31 | 0.000 | 66 | -18 | 26 |
| | L | 7.34 | 0.040 | - part of Cluster *c* - | | -58 | -30 | 26 |
| Superior parietal lobule | L | 8.98 | 0.005 | 32 | 0.000 | -28 | -48 | 56 |
| Broca's Area | L | 8.92 | 0.005 | 14 | 0.003 | -52 | 2 | 34 |
| **Cerebellar Regions** | | | | | | | | |
| CB V | R | 7.78 | 0.022 | 30 | 0.000 | 18 | -50 | -26 |
| CB VI | L | 7.61 | 0.027 | 10 | 0.005 | -34 | -52 | -20 |
| **Subcortical Regions** | | | | | | | | |
| Putamen | L | 14.94 | 0.000 | 1439[d] | 0.000 | -8 | -10 | 0 |
| | R | 12.58 | 0.000 | 1104[e] | 0.000 | 30 | -4 | 10 |
| | L | 12.16 | 0.000 | - part of Cluster *d* - | | -32 | -12 | -2 |
| | R | 11.90 | 0.000 | - part of Cluster *e* - | | 26 | -2 | 0 |

Superscripts on the cluster volume, when present, refer to clusters with multiple peaks. Coordinates are expressed in MNI space.

### Supplementary Figures and Tables

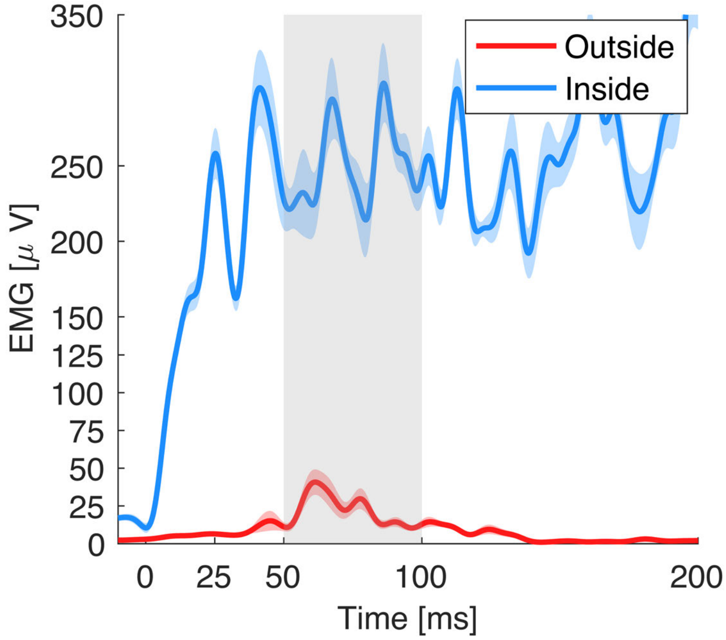


**Figure S5.** Stretch-evoked EMG signal measured using off-the-shelf electrodes, amplifier, and signal processing pipeline in response to a 125 deg/s RaH perturbation for the Flexor Carpi Radialis (FCR) for a representative subject.

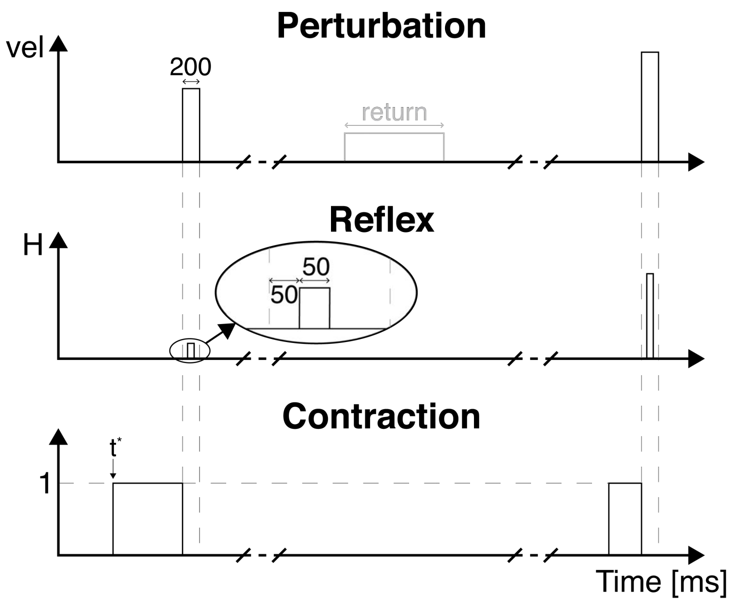


**Figure S6.** Graphical representation of the regressors used in the statistical analysis of the fMRI data. The value $t^\star$ in the contraction regressor represents the time at which the pre-condition isometric torque increased above 25% of the desired background torque.

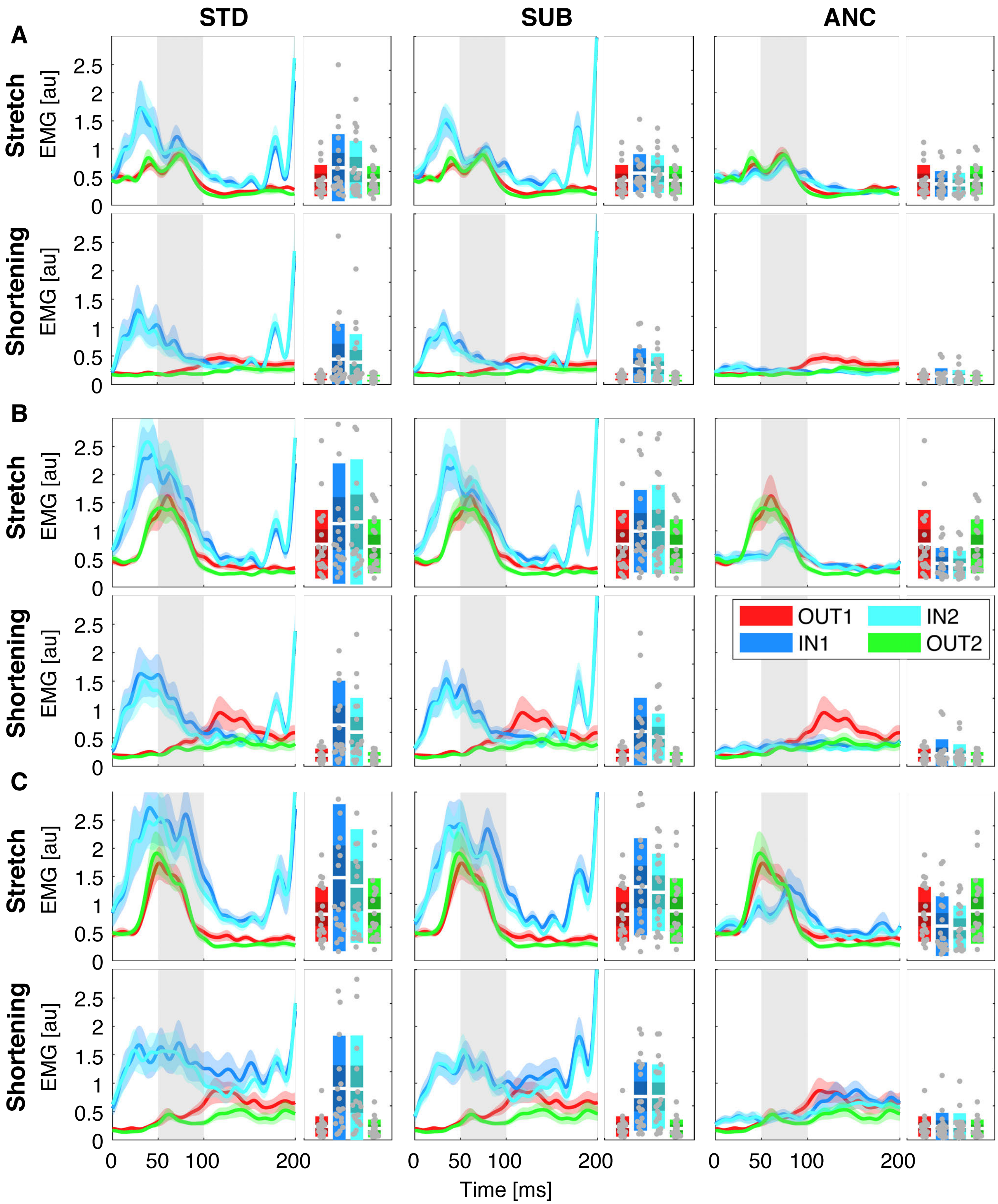


**Figure S7.** Group-average of the stretch-evoked EMG signal measured using different processing pipelines in response to a (**A**) 50 deg/s, (**B**) 125 deg/s, and (**C**) 200 deg/s RaH perturbation for both muscles undergoing stretch or shortening for experiment 1. Plots are obtained with a combination of the novel data collection hardware with a standard signal processing pipeline (left), a time-domain subtraction of measured and reference signals (center), and the novel adaptive noise cancellation pipeline (right). In all graph the thick line represents the mean and the shaded area the 95% confidence interval. To facilitate interpretation on the graphs a gray shaded area has been included in each graph in the time interval where a LLR is expected.

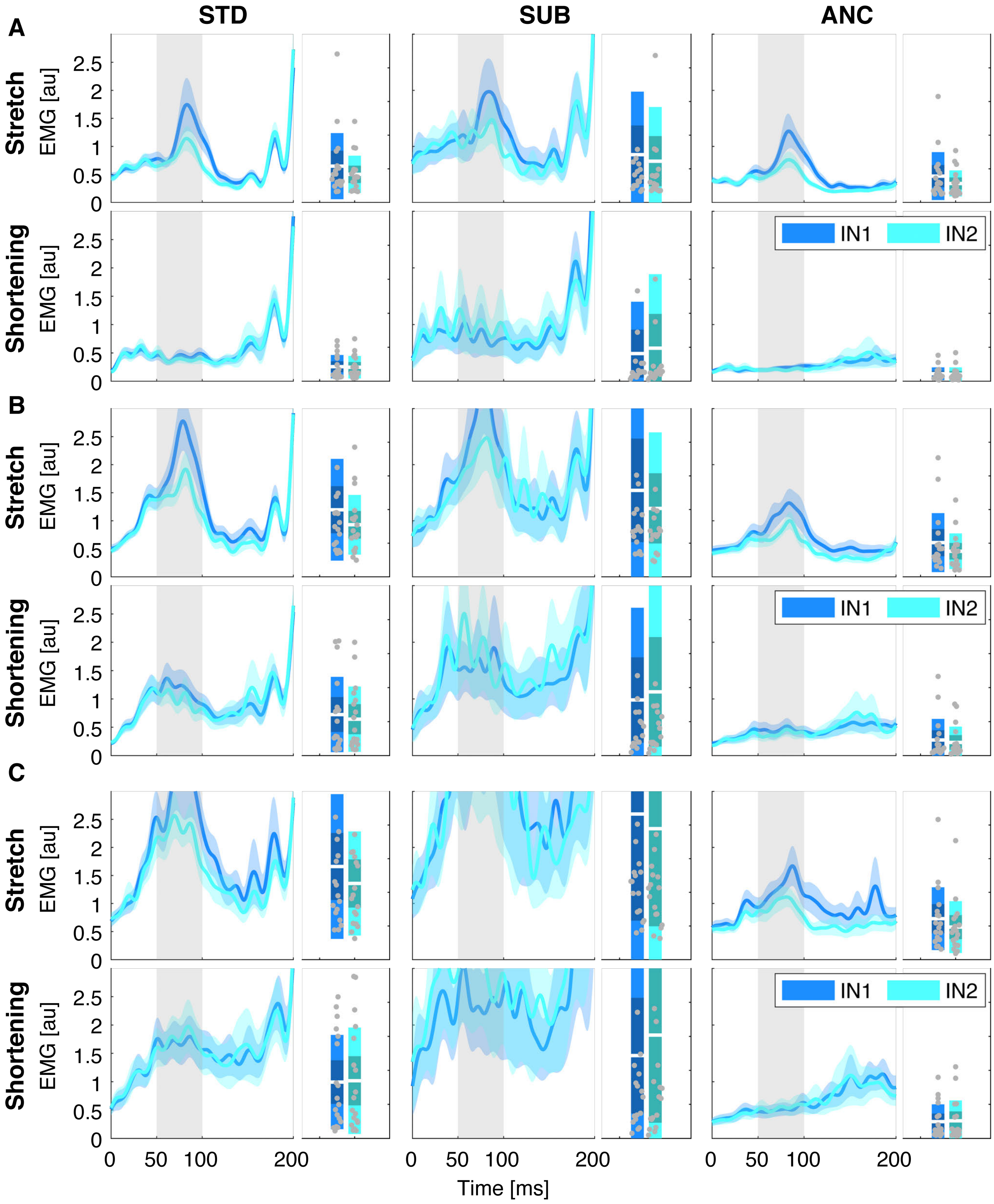


**Figure S8.** Group-average of the stretch-evoked EMG signal measured using different processing pipelines in response to a (**A**) 50 deg/s, (**B**) 125 deg/s, and (**C**) 200 deg/s RaH perturbation for both muscles undergoing stretch or shortening for experiment 2. Plots are obtained with a combination of the novel data collection hardware with a standard signal processing pipeline (left), a time-domain subtraction of measured and reference signals (center), and the novel adaptive noise cancellation pipeline (right). In all graph the thick line represents the mean and the shaded area the 95% confidence interval. To facilitate interpretation on the graphs a gray shaded area has been included in each graph in the time interval where a LLR is expected.

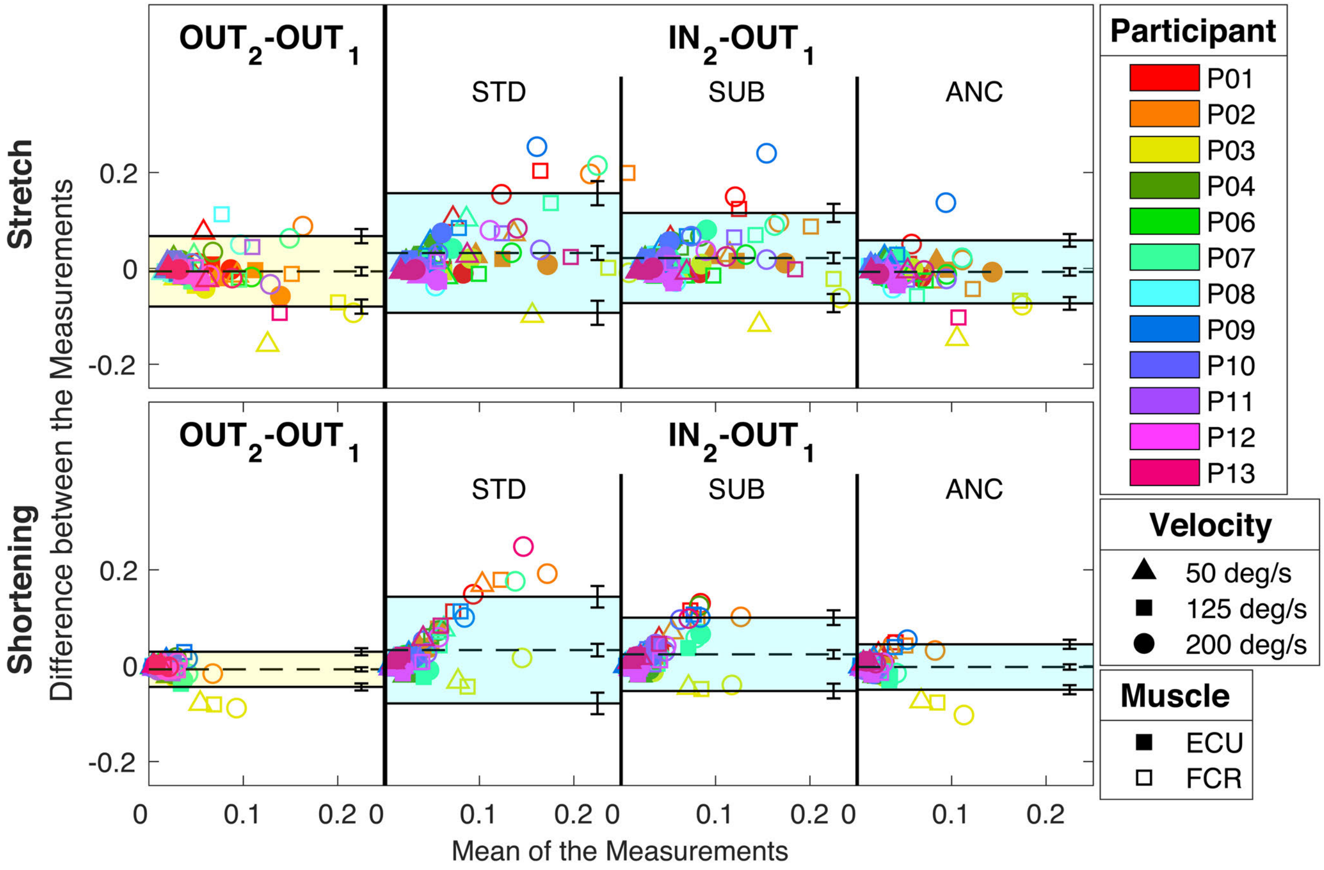


**Figure S9.** Bland-Altman plots for the comparisons $OUT_2$ vs. $OUT_1$ (on the left) and $IN_2$ vs. $OUT_1$ when different filtering pipelines are considered (on the right). Datapoints in each of the plots model participants with colors, perturbation velocities with shapes, and muscles from which the EMG is recorded with fillings. The shaded area represents the interval between the negative and positive limits of agreement, while the dashed line represent the bias. Error bars on the side of each plot represent the 95% confidence intervals by which each of the parameter is estimated. Top and bottom plots distinguish the muscle activity measured in response of the two stimulus directions, respectively stretch on top and shortening on the bottom.

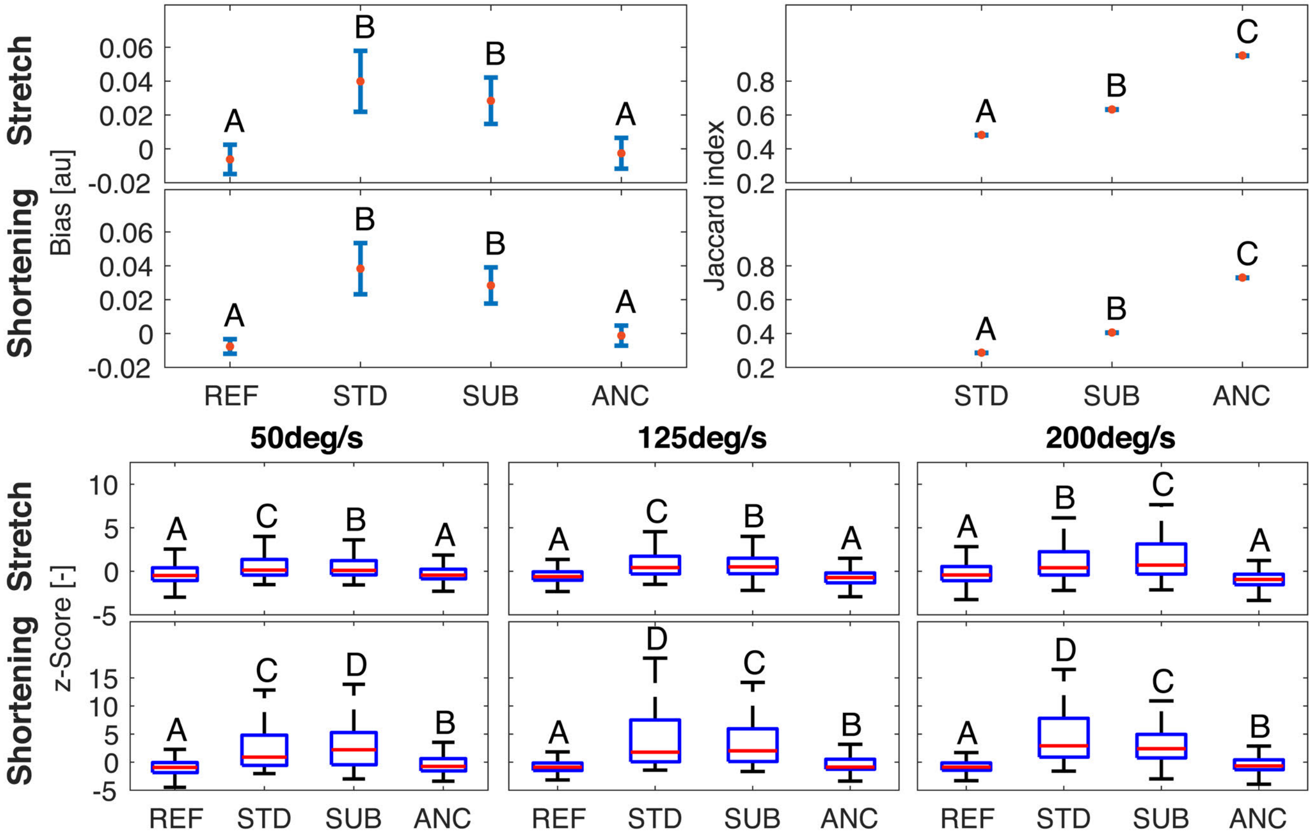


**Figure S10.** Mean and 95% confidence intervals for the bias (**A**), and the Jaccard index (**B**) obtained for the different filtering pipelines. Test-retest parameters measured for the two stimulus directions are reported in different rows. Letters indicate rankings of the test-retest parameters. Bars that do not share a letter have a statistically different mean. (**C**) Box plots representing the distributions of standardized *z*-scores measured for different combinations of perturbation velocity and filtering pipeline. Distributions obtained for the two stimulus directions are reported in different rows. Letters indicate rankings of the variance of each distribution; boxes that do not share a letter have a statistically different variance.

**Table S8.** Results of the Bland-Altman analysis for session $IN_1$

| | **$OUT_2$-$OUT_1$** | | **$IN_1$-$OUT_1$** STD | | SUB | | ANC | |
|---|---|---|---|---|---|---|---|---|
| | Stretch | Shortening | Stretch | Shortening | Stretch | Shortening | Stretch | Shortening |
| Bias | -0.007 | -0.007 | 0.032 | 0.030 | 0.023 | 0.021 | -0.001 | 0.002 |
| 95% CI | [-0.015, 0.001] | [-0.012, 0.003] | [0.017, 0.047] | [0.017 0.043] | [0.010, 0.035] | [0.012, 0.030] | [-0.009, 0.007] | [-0.007, 0.003] |
| LoA- | -0.070 | -0.043 | -0.096 | -0.080 | -0.082 | -0.056 | -0.072 | -0.046 |
| 95% CI | [-0.082, -0.057] | [-0.050, -0.036] | [-0.123, -0.071] | [-0.102, -0.058] | [-0.103, -0.061] | [-0.072, -0.040] | [-0.087, -0.058] | [-0.055, -0.037] |
| LoA+ | 0.056 | 0.028 | 0.161 | 0.140 | 0.128 | 0.099 | 0.070 | 0.042 |
| 95% CI | [0.043, 0.068] | [0.021, 0.035] | [0.135, 0.187] | [0.118, 0.162] | [0.107, 0.149] | [0.084, 0.115] | [0.056, 0.085] | [0.033, 0.051] |
| Jaccard | | | 0.489 | 0.324 | 0.593 | 0.457 | 0.843 | 0.783 |
| 95% CI | | | [0.4890 0.4896] | [0.3241, 03245] | [0.5930, 0.5937] | [0.4562, 0.4567] | [0.8424, 0.8432] | [0.7828 0.7836] |

**Table S9.** Standard deviation for the set of normalized scores obtained for session $IN_1$

| | vel [deg/s] | REF | STD | SUB | ANC |
|---|---|---|---|---|---|
| Stretch | 50 | 0.896 | 1.353 | 1.256 | 0.905 |
| | 125 | 0.858 | 1.420 | 1.285 | 0.933 |
| | 200 | 1.175 | 2.287 | 2.118 | 1.021 |
| Shortening | 50 | 1.209 | 3.644 | 3.376 | 1.161 |
| | 125 | 1.051 | 4.980 | 3.789 | 1.434 |
| | 200 | 1.143 | 4.020 | 3.587 | 1.339 |

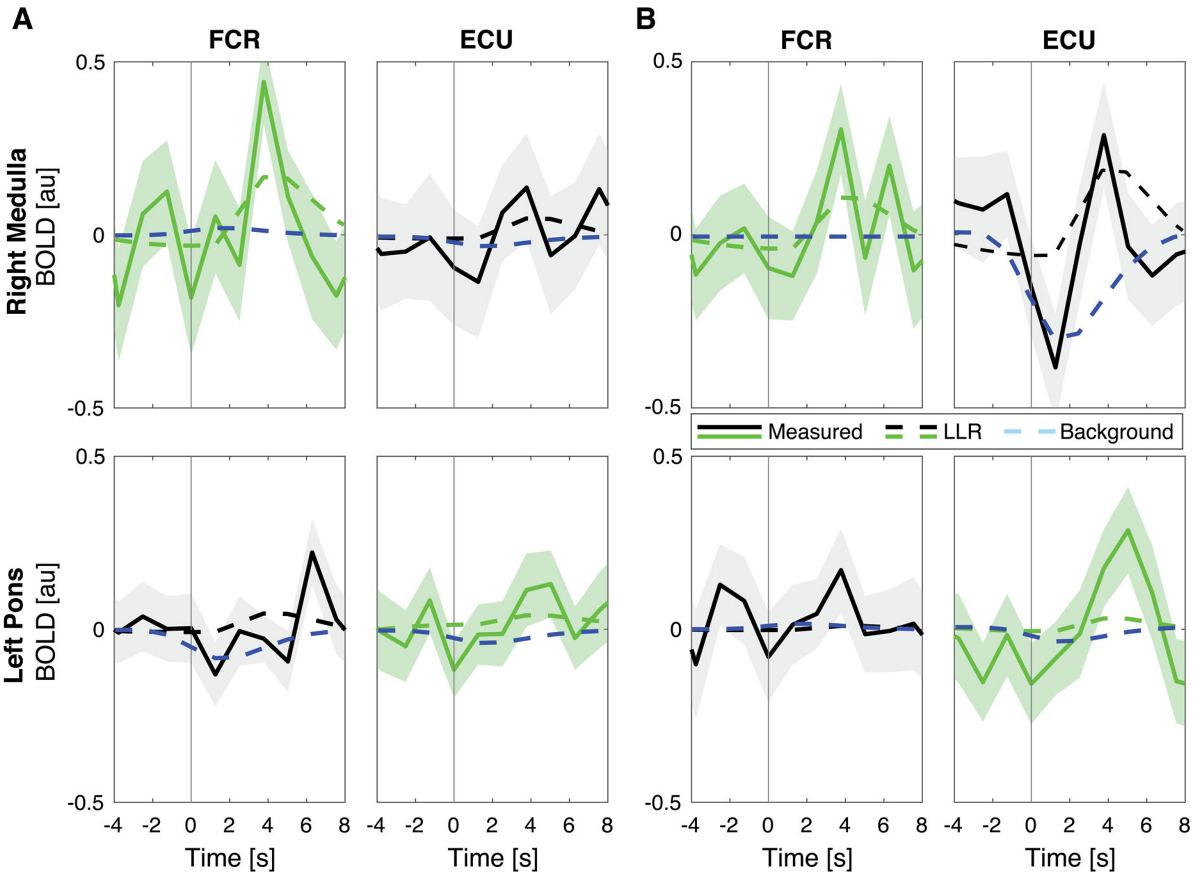


**Figure S11.** Average BOLD signal measured in response to muscle stretch in the selected voxels, for the selected participants. Specifically, **A** include the data that refer to the subjects that showed the greatest difference between the statistical scores specific to the FCR and ECU regressors, while **B** include the data that refer to those subject that showed similar $t$ score for the two regressors (Fig. 5). In all graphs, the solid line shows the average residual BOLD response after variance associated to nuisance regressors is removed, while the shaded area represents the 95% confidence interval of the mean response. Dashed lines show the average modeled response respectively for the LLR-specific regressors (green or black line), and for the background-specific regressor (light blue line). Modeled response for the LLR-specific regressor was scaled based on the appropriate $\beta$ coefficient estimated by the GLM model, while modeled background response, which is not included in the primary GLM, was arbitrarily scaled to match the same amplitude of the LLR-specific regressor. Measured and LLR-specific signals are then color-coded to distinguish the response that is significant at the group level for that specific muscle (green), from the one that is not significant (black).

**Table S10.** Results of the Bland-Altman analysis for session $IN_2$

| | **$OUT_2$-$OUT_1$** | | **$IN_2$-$OUT_1$** STD | | SUB | | ANC | |
|---|---|---|---|---|---|---|---|---|
| | Stretch | Shortening | Stretch | Shortening | Stretch | Shortening | Stretch | Shortening |
| Bias | -0.006 | -0.008 | 0.032 | 0.033 | 0.022 | 0.024 | -0.007 | 0.003 |
| 95% CI | [-0.015, 0.002] | [-0.012, 0.003] | [0.017, 0.047] | [0.019 0.047] | [0.011, 0.033] | [0.015, 0.033] | [-0.015, 0.001] | [-0.008, 0.003] |
| LoA- | -0.079 | -0.044 | -0.093 | -0.079 | -0.072 | -0.053 | -0.073 | -0.050 |
| 95% CI | [-0.094, -0.064] | [-0.050, -0.037] | [-0.118, -0.068] | [-0.101, -0.056] | [-0.091, -0.053] | [-0.068, -0.037] | [-0.086, -0.060] | [-0.059, -0.040] |
| LoA+ | 0.067 | 0.029 | 0.157 | 0.144 | 0.116 | 0.100 | 0.058 | 0.044 |
| 95% CI | [0.052, 0.082] | [0.022, 0.037] | [0.132, 0.182] | [0.121, 0.166] | [0.097, 0.135] | [0.085, 0.116] | [0.045, 0.072] | [0.035, 0.054] |
| Jaccard | | | 0.589 | 0.330 | 0.715 | 0.480 | 0.895 | 0.775 |
| 95% CI | | | [0.5875 0.5901] | [0.3290, 03311] | [0.7136, 0.7163] | [0.4786, 0.4810] | [0.8933, 0.8959] | [0.7740 0.7766] |

**Table S11.** Standard deviation for the set of normalized scores obtained for session $IN_2$

| | vel [deg/s] | REF | STD | SUB | ANC |
|---|---|---|---|---|---|
| Stretch | 50 | 1.042 | 1.386 | 1.326 | 0.945 |
| | 125 | 0.835 | 1.476 | 1.375 | 0.910 |
| | 200 | 1.181 | 2.158 | 2.230 | 0.993 |
| Shortening | 50 | 1.299 | 4.173 | 3.832 | 1.562 |
| | 125 | 1.117 | 5.732 | 3.892 | 1.411 |
| | 200 | 1.131 | 4.664 | 3.722 | 1.469 |

**Table S12.** Test-retest reliability of fMRI data

$H_{FCR}$

| | | Cortical ROIs | | | | | | | | | |
|---|---|---|---|---|---|---|---|---|---|---|---|
| | | M1 | | PM | | S1 | | SPL | | IPS | |
| | | R | **L** | R | **L** | R | **L** | R | **L** | R | **L** |
| ICC | medICC | 0.57 (0.01) | 0.57 (0.01) | 0.54 (0.01) | 0.56 (0.01) | 0.55 (0.00) | 0.71 (0.01) | 0.63 (0.00) | 0.55 (0.00) | 0.56 (0.03) | 0.39 (0.02) |
| | maxICC | 0.88 | 0.86 | 0.83 | 0.88 | 0.87 | 0.94 | 0.9 | 0.87 | 0.72 | 0.74 |
| | ICCv | 0.50 (0.01) | 0.69 (0.01) | 0.59 (0.01) | 0.74 (0.01) | 0.57 (0.01) | 0.76 (0.01) | 0.53 (0.02) | 0.55 (0.01) | 0.48 (0.02) | 0.48 (0.02) |
| | Sg | 0 | 0.71 | 0.06 | 0.6 | 0 | 0.61 | 0.1 | 0.24 | 1 | 0.58 |
| | Ss | 0.19 (0.01) | 0.44 (0.02) | 0.35 (0.02) | 0.43 (0.02) | 0.28 (0.01) | 0.64 (0.01) | 0.32 (0.02) | 0.26 (0.02) | 0.33 (0.03) | 0.23 (0.02) |

| | | Cortical ROIs | | Subcortical ROIs | | | | | Whole Brain |
|---|---|---|---|---|---|---|---|---|---|
| | | Ce | | Th | | Pt | | Bs | |
| | | **R** | L | R | **L** | R | **L** | | |
| ICC | medICC | 0.61 (0.00) | 0.58 (0.00) | 0.37 (0.01) | 0.43 (0.00) | 0.43 (0.01) | 0.49 (0.01) | 0.28 (0.01) | 0.52 (0.00) |
| | maxICC | 0.9 | 0.96 | 0.81 | 0.72 | 0.76 | 0.81 | 0.98 | 1 |
| | ICCv | 0.61 (0.01) | 0.60 (0.01) | 0.44 (0.01) | 0.38 (0.01) | 0.32 (0.01) | 0.32 (0.01) | 0.31 (0.01) | 0.62 (0.01) |
| | Sg | 0.86 | 0.84 | 0.4 | 0.52 | 0.45 | 0.57 | 0.06 | 0.5 |
| | Ss | 0.42 (0.02) | 0.46 (0.01) | 0.13 (0.01) | 0.13 (0.01) | 0.06 (0.01) | 0.14 (0.01) | 0.06 (0.01) | 0.44 (0.01) |

$H_{ECU}$

| | | Cortical ROIs | | | | | | | | | |
|---|---|---|---|---|---|---|---|---|---|---|---|
| | | M1 | | PM | | S1 | | SPL | | IPS | |
| | | R | **L** | R | **L** | R | **L** | R | **L** | R | **L** |
| ICC | medICC | 0.51 (0.01) | 0.57 (0.00) | 0.37 (0.00) | 0.47 (0.00) | 0.49 (0.00) | 0.58 (0.01) | 0.58 (0.00) | 0.54 (0.00) | 0.40 (0.01) | 0.39 (0.01) |
| | maxICC | 0.73 | 0.85 | 0.74 | 0.74 | 0.79 | 0.84 | 0.82 | 0.77 | 0.52 | 0.58 |
| | ICCv | 0.48 (0.01) | 0.69 (0.01) | 0.56 (0.01) | 0.72 (0.01) | 0.65 (0.01) | 0.75 (0.01) | 0.61 (0.01) | 0.64 (0.01) | 0.47 (0.02) | 0.41 (0.02) |
| | Sg | 0 | 0.45 | 0.14 | 0.23 | 0.12 | 0.58 | 0.01 | 0.08 | NaN | 0.18 |
| | Ss | 0.16 (0.01) | 0.43 (0.01) | 0.26 (0.01) | 0.46 (0.02) | 0.27 (0.01) | 0.59 (0.02) | 0.26 (0.02) | 0.37 (0.02) | 0.10 (0.01) | 0.17 (0.02) |

| | | Cortical ROIs | | Subcortical ROIs | | | | | Whole Brain |
|---|---|---|---|---|---|---|---|---|---|
| | | Ce | | Th | | Pt | | Bs | |
| | | **R** | L | R | **L** | R | **L** | | |
| ICC | medICC | 0.50 (0.00) | 0.34 (0.00) | 0.39 (0.01) | 0.37 (0.00) | 0.55 (0.01) | 0.54 (0.01) | 0.24 (0.00) | 0.48 (0.00) |
| | maxICC | 0.88 | 0.95 | 0.79 | 0.72 | 0.81 | 0.74 | 0.94 | 1 |
| | ICCv | 0.58 (0.01) | 0.53 (0.02) | 0.46 (0.01) | 0.41 (0.02) | 0.30 (0.02) | 0.34 (0.01) | 0.29 (0.01) | 0.62 (0.01) |
| | Sg | 0.75 | 0.74 | 0.74 | 0.6 | 0.14 | 0.41 | 0.12 | 0.41 |
| | Ss | 0.39 (0.02) | 0.30 (0.01) | 0.13 (0.01) | 0.12 (0.01) | 0.18 (0.02) | 0.14 (0.01) | 0.06 (0.01) | 0.41 (0.01) |
